# HalluDesign: Protein Optimization and de novo Design via Iterative Structure Hallucination and Sequence Design

**DOI:** 10.1101/2025.11.08.686881

**Authors:** Minchao Fang, Chentong Wang, Jungang Shi, Fengbai Lian, Qihan Jin, Zhe Wang, Yanzhe Zhang, Peipei Chen, Zhanyuan Cui, Yanjun Wang, Ze Zhang, Yitao Ke, Qingzheng Han, Longxing Cao

**Author notes:** These authors contributed equally to this work (Minchao Fang, Chentong Wang, Jungang Shi, Fengbai Lian).

## Abstract

Deep learning has revolutionized biomolecular modeling, enabling the prediction of diverse structures with atomic accuracy. However, leveraging the atomic-level precision of the structure prediction model for de novo design remains challenging. Here, we present HalluDesign, a general all-atom framework for protein optimization and de novo design, which iteratively update protein structure and sequence. HalluDesign harnesses the inherent hallucination capabilities of AlphaFold3-style structure prediction models and enables fine-tune free, forward-pass only sequence-structure co-optimization. Structure conditioning at different noise level in the structure prediction stage allows precise control over the sampling space, facilitating tasks from local and global protein optimization to de novo design. We demonstrate the versatility of this framework by optimizing suboptimal structures, rescuing previously unsuccessful designs, designing new biomolecular interactions and generating new protein structures from scratch. Experimental characterization of a diverse set of proteins—including protein binders for small molecules, a metal ion and proteins; antibody design of phosphorylation-specific peptide; and monomeric proteins—revealed high design success rates and excellent structural accuracy. Together, our comprehensive computational and experimental results highlight the broad utility of this framework. We anticipate that HalluDesign will further unlock the modeling and design potential of AlphaFold3-like models, enabling the robust creation of complex biomolecules for a wide range of biotechnological applications.

## Main Text

Deep learning–based neural networks for protein structure prediction have revolutionized biological research field, enabling computational modeling of protein structures with atomic accuracy primarily based on sequence information (*1–4*). Nevertheless, translating the atomic-level precision of these state-of-the-art models into a robust framework for de novo design—rather than just structure prediction—remains a significant challenge. Recent progress in protein design has leveraged the pretrained structure prediction models. Early efforts employed intensive Monte Carlo sampling in the discrete sequence space (*5, 6*), starting from random sequences and utilizing structure prediction after single mutations to guide the sampling trajectory. Current approaches can be broadly classified into two categories: gradient-based backpropagation methods and fine-tuning strategies. Gradient-based methods iteratively optimize sequence and structure compatibility by minimizing network loss (*7–10*); however, they typically require high computational costs and often struggle to converge due to the ruggedness of the sequence–structure landscape. In contrast, the fine-tuning approaches retrain structure prediction models for protein backbone generation through a generative diffusion process (*11–15*); however, they typically require generating a large number of scaffolds and subsequently performing extensive sequence sampling to identify candidates that can pass in silico structure prediction filters, such as AlphaFold2 or AlphaFold3, before proceeding to experimental validation.

Recently, AlphaFold3 has expended the scope of structure prediction beyond proteins to include small molecules, nucleic acids, ions, and covalent modifications, achieving unprecedented accuracy (*16, 17*). This breakthrough, along with the various open-source reproductions (*18–20*), has spurred a new wave of adaptation and fine-tuning for the modeling and design of diverse biomolecules (*21–24*). The AlphaFold3 architecture is primarily composed of a pairformer module, which extracts residue/atom-level and pairwise representations from input features, and a diffusion module, which generates all-atom structures from Gaussian noise conditioned on the information provided by the pairformer. The generative diffusion process inherently results in “hallucination” —the generation of plausible-looking structures in the low-confidence regions (**Fig. 1A**) (*16, 17*). While this property is an undesirable side-effect for structure prediction, we hypothesized that it could be repurposed for structure optimization and de novo design, without requiring model fine-tuning or computationally intensive loss backpropagation. Here, we introduce HalluDesign—a hallucination-driven, forward-pass-only framework that leverages the AlphaFold3-style structure prediction models to iteratively optimize and design both protein sequences and structures.

**Fig. 1.**
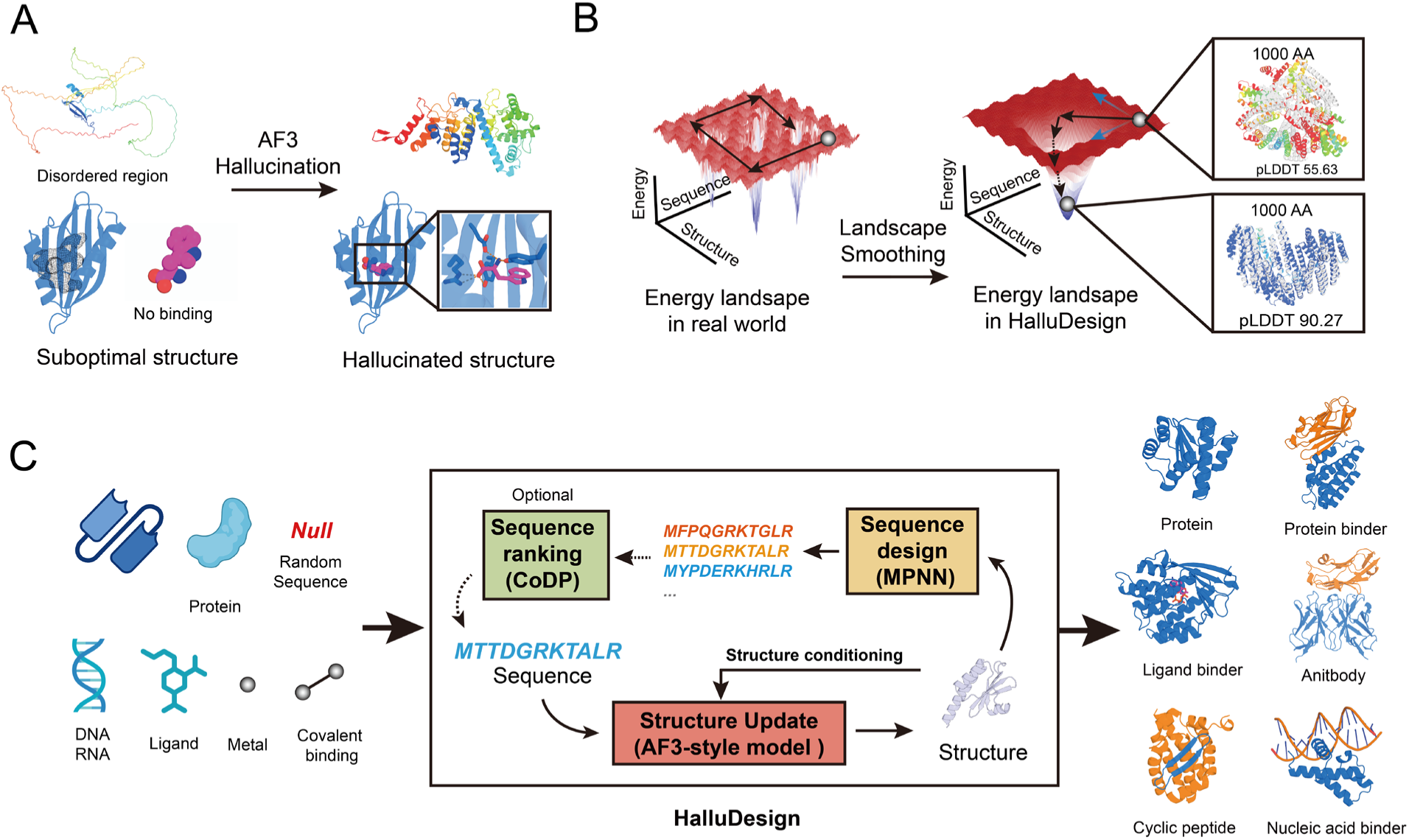
Overview of the HalluDesign framework. **(A)** AlphaFold3-like models leverage the inherent ‘hallucination’ capability of the diffusion structure module to generate plausible structures even in low-confidence regions. **(B)** The hallucination effect smooths the rugged sequence–structure landscape, allowing efficient protein optimization via iterative sequence–structure sampling. **(C)** Schematic representation of the HalluDesign workflow. HalluDesign employs alternating rounds of sequence and structure updates for structure-sequence co-optimization. The CoDP model can be optionally applied to select the most compatible sequence at each iteration to accelerate convergence. By leveraging advanced biomolecular modeling, HalluDesign enables the design and optimization of diverse biomolecule structures, including proteins, DNA, RNA, and small molecules.

### HalluDesign methodology

The real protein sequence–structure landscape is highly rugged (*25–28*), making it challenging to find optimal solutions through iterative traversing the sequence and structure space. We reasoned that diffusion-based structure prediction models intrinsically smooth this rugged landscape, which is often observed as ‘hallucination’ effect (**Fig. 1B**). To leverage this property for optimization and de novo design, we developed HalluDesign, an iterative framework designed to co-optimize and co-design protein sequence and structure (**Fig. 1C**), alternating between a structure update stage and a sequence design stage.

This iterative framework begins with the structure update stage, where the diffusion module of an AlphaFold3-style model leverages its ‘hallucination’ capabilities to refine the protein structure, guiding it towards a more optimal conformation conditioned on the input sequence. This process enables the generation of plausible-looking conformations even for suboptimal protein sequences lacking stable folds and interactions. In the subsequent sequence design stage, an inverse folding deep learning model generates a new amino acid sequence that is more compatible with the updated structure. This new sequence is then fed back into the next structure update iteration, serving to better condition the diffusion module and ensure the model operates with increasingly compatible sequence and structure pair. Notably, the entire process requires only forward passes through the structure prediction and sequence design models, requiring neither model fine-tuning nor computationally intensive loss backpropagation.

The structure update stage provides a unified and controllable strategy by utilizing a truncated diffusion approach. Rather than initializing the diffusion process of the AlphaFold3 structure module with random Gaussian noise, a user-defined noise level is applied to the initial or a previous iteration’s structure (**Fig. 2A** and **fig. S1A**). This noised structure then serves as the input for the reverse diffusion process to generate an updated structure. This method offers precise control over the structural exploration by truncating the diffusion trajectory at various timesteps (**Fig. 2B**). Shorter trajectories, corresponding to lower noise levels, prompt local refinement, which is well-suited for optimization tasks requiring the preservation of the initial topology. In contrast, longer trajectories, corresponding to higher noise levels, enable broader exploration of structure landscape to escape from local minima.

**Fig. 2.**
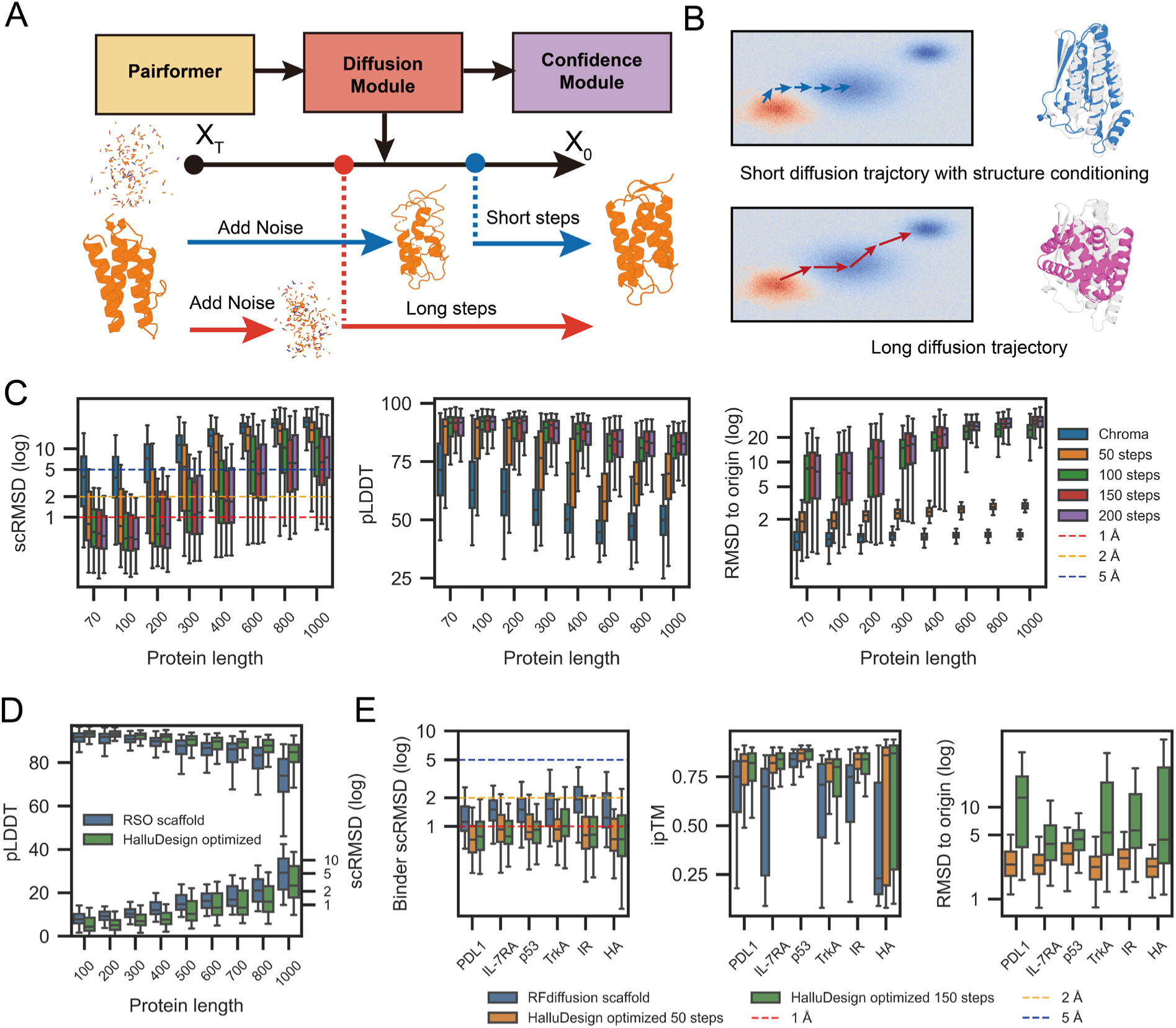
HalluDesign for structure optimization. **(A)** Truncating the diffusion trajectory at different steps and applying prior-structure conditioning allows precise control over the extent of structural exploration. **(B)** Short trajectories prioritize local refinement and topology preservation, while long trajectories enable broader conformation exploration to escape local minima. **(C)** HalluDesign optimization of Chroma-generated monomers. Seven optimization cycles were conducted, with truncated diffusion trajectories of 50, 100, 150 and 200 steps, respectively. The computational metrics (evaluated by ESMFold) include self-consistency RMSD (scRMSD), pLDDT and structural deviation from the initial model. **(D)** HalluDesign optimization of RSO-generated structures (ESMFold as the orthogonal evaluator). **(E)** HalluDesign optimization of RFdiffusion-generated protein binders. The impact of short (50 steps) versus long (150 steps) diffusion trajectories is assessed via binder scRMSD (aligned by target), interface pTM (ipTM), and structural deviation from the initial design across different targets (evaluated by AlphaFold2-Multimer).

The sequence design stage fulfills a dual role by determining the final amino acid sequence and, at each intermediate iteration, providing a backbone-compatible sequence to optimally condition the subsequent structure update. This is achieved using inverse folding models such as ProteinMPNN and LigandMPNN (*29, 30*), which allow efficient sampling of candidate sequences for the current backbone. We hypothesized that selecting candidate sequence with higher structural compatibility, rather than choosing one at random, would facilitate more rapid convergence. Thus, we developed CoDP (Contrastive-learning-based Distogram Prediction Model) (**fig. S1B** and **fig. S2**), a lightweight model that rapidly ranks candidate sequences via distogram prediction (*1, 3, 4*) and contrastive learning (*31, 32*) (see more in Supplementary Methods). To ensure rigorous benchmarking of HalluDesign’s core optimization logic, CoDP is excluded from results unless otherwise noted, and a single sequence is instead randomly selected at each step.

### HalluDesign for protein optimization

While state-of-the-art generative models typically use AlphaFold2/3 as an external ‘oracle’ to select high-quality candidates from a large set of designs (*13*), we hypothesized that HalluDesign could instead actively optimize these existing designs. We first applied HalluDesign to optimize monomeric scaffolds generated by Chroma (*33*). Both structural and confidence metrics improved steadily across iterative cycles assessed via an orthogonal model ESMFold (*4*) (**Fig. 2C** and **fig. S3**). This optimization performance was further amplified by employing CoDP for sequence selection, which consistently yielded superior structural confidence and accuracy compared to random sampling (**fig. S4**). Notably, the intrinsic confidence metrics of the HalluDesign foundation model showed strong correlations with those obtained from the orthogonal evaluation model (**fig. S5**). Moreover, we observed distinct optimization behaviors by varying the length of the diffusion trajectory: shorter trajectories (e.g., 50 steps) preserved the input topology but converged more slowly, making them ideal for refinement. Longer trajectories (100–200 steps) explored broader structural landscapes and reached high-confidence solutions rapidly, typically within 6–7 cycles (**fig. S3**). This trade-off was particularly pronounced for large proteins (>300 aa), which required longer trajectories or additional cycles to accommodate the substantial conformational changes needed for convergence.

To further assess the generalizability of HalluDesign’s optimization capabilities, we applied it to scaffolds generated by RSO (*34*), a state-of-the-art design method based on AlphaFold2 backpropagation. We selected RSO as a challenging benchmark as it is one of the most performant approaches for monomer generation, particularly for large proteins. Remarkably, our forward-pass-only approach was able to further improve computational metrics for the RSO backbones evaluated by ESMFold (*4*) (**Fig. 2D** and **fig. S6**). This finding suggests that the protein sequence-structure landscape modeled by HalluDesign is smoother and more amenable to optimization than the AF2 landscape explored by gradient-based methods. In contrast, we observed that RFdiffusion partial diffusion failed to refine these same RSO backbones (*14, 35*). This highlights that protein structure landscape is more accurately modeled by HalluDesign than that of RFdiffusion, underscoring the necessity of leveraging HalluDesign as the foundation platform for optimization.

Having demonstrated HalluDesign’s efficacy on monomeric proteins, we next evaluated its ability to refine biomolecular interactions. We applied HalluDesign to protein binders generated by RFdiffusion from their published PPI supplementary dataset (*35*). These binders exhibited high computational metrics and have been experimentally characterized. Remarkably, HalluDesign further improved both structural accuracy and confidence metrics across all targets evaluated via Alphafold2-multimer (*2*) (**Fig. 2E** and **fig. S7**). We found that structure conditioning with shorter diffusion trajectories achieved comparable optimization performance to longer ones, while better preserving structural consistency with the input model.

Then we leveraged the all-atom capabilities of the underlying AF3 model to optimize small-molecule binders. We curated a diverse set of small-molecule ligands spanning a broad range of sizes and chemical properties (**fig. S8** and **table S1**). We then generated initial binders for these ligands using the RFdiffusion all-atom (*36*) and the BoltzDesign1 (*21*). While these initial designs showed modest quality, optimization by HalluDesign yielded significant improvements in computational metrics across all targets and diffusion trajectory lengths, regardless of the source of the starting structure (**fig. S9** and **fig. S10)**. Interestingly, while CoDP ranks sequences based solely on protein sequence-structure compatibility, the resulting enhancement in monomer quality effectively translated into improved ligand binding prediction metrics (**fig. S4E**). Crucially, these enhancements were consistently observed when assessed by other independent all-atom prediction models (*18–20*) (**fig. S11** and **fig. S12)**. This cross-model validation demonstrates that HalluDesign achieves physically meaningful improvements in interaction quality, rather than model-specific artifacts or overfitting to a particular foundation model. We attribute this success to the enhanced intermolecular prediction capabilities inherent to the AF3 architecture, as well as HalluDesign allows to simultaneously co-optimize both the protein structure and the ligand conformation—a distinct advantage over the fixed-ligand design approaches (*36*).

### HalluDesign for scaffold-based de novo protein design

Beyond refining existing structures, we hypothesized that HalluDesign could design new interaction with targets using existing protein scaffolds that do not naturally interact with those targets. To test this capability, we performed de novo design of small molecule binders on two distinct protein topologies. First, we employed the nuclear transport factor 2 (NTF2) scaffold, a compact and highly engineerable fold known to accommodate diverse ligands (*10, 37*). Second, to verify the method generalizability to other protein topologies, we generated a large set of pseudocycle scaffolds with pockets aligned along their pseudo-symmetric axis (see more in the supplementary methods) (*38*). For both scaffold types, we adopted a “black hole initialization” strategy—placing a random ligand conformer at the centre of mass of the protein (**Fig. 3A**). We then applied HalluDesign to generate high-quality protein–ligand complex structures. We truncated the diffusion step to 150 steps unless otherwise specified for design, as this configuration provided similar optimization performance to the default full diffusion trajectory (200 steps) (**fig. S3** and **fig. S10**).

**Fig. 3.**
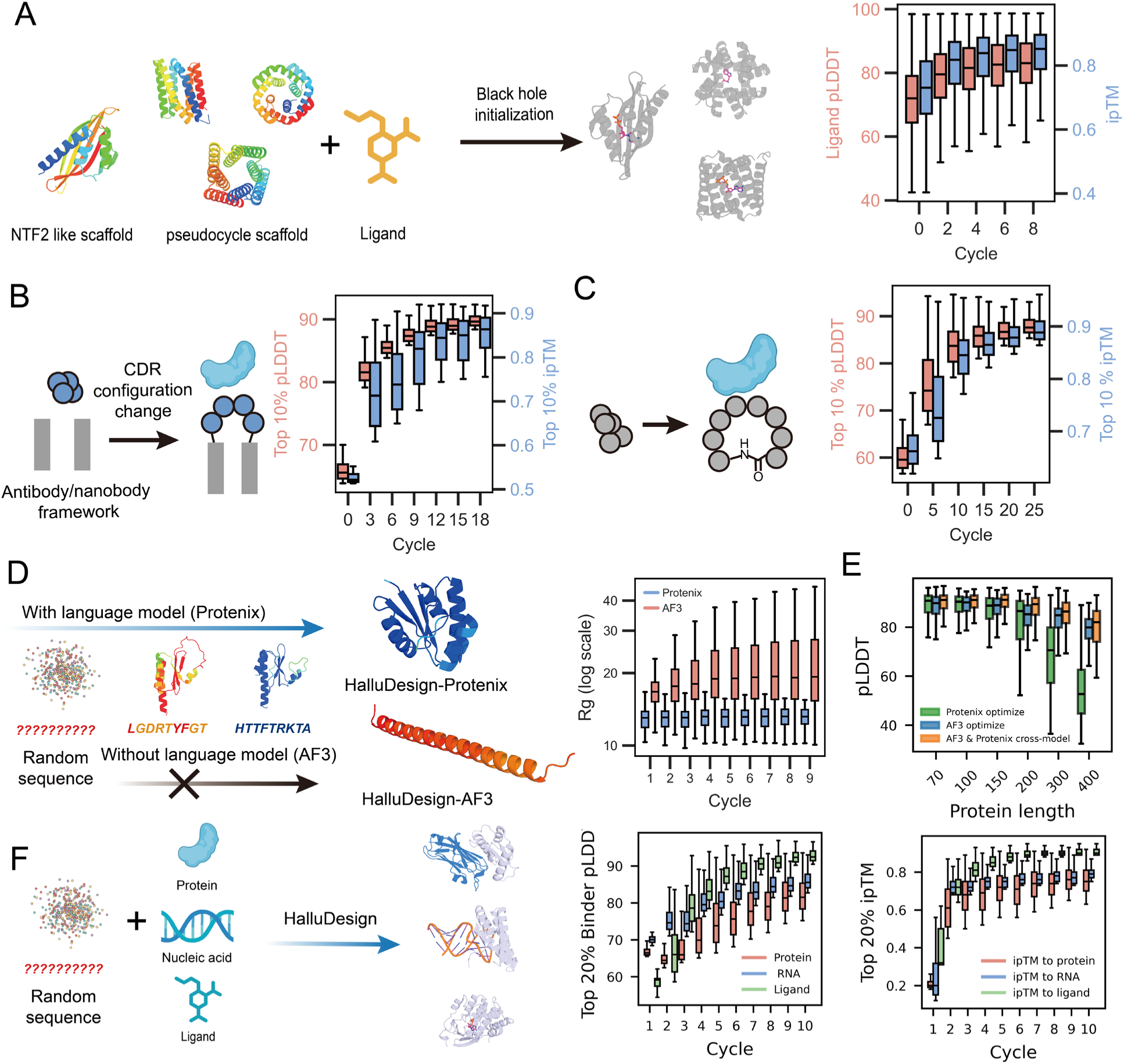
HalluDesign for de novo protein design. **(A)** HalluDesign for small molecule binder design using pocket-containing scaffolds. Starting from a ligand randomly positioned at the center of a protein scaffold, HalluDesign enables de novo generation of protein–ligand complexes. The average computational metrics for the in silico small molecule dataset (**fig. S8**), as assessed by the AF3 confidence metrics, are reported across design cycles. **(B)** HalluDesign for antibody design. PDL1 target-specific antibodies are generated de novo, initiating from randomized CDR loop configurations. The computational metrics (from AF3) of the top 10% highest-scoring designs are reported (**fig. S15C**). **(C)** HalluDesign for cyclic peptide binder design. Starting from random cyclic peptide sequence, HalluDesign enables the generation of target specific cyclic peptides. Average computational metrics for the top 10% highest-scoring designs for four protein targets (PDL1, TNFα receptor, MCL1, and EGFR) consistently demonstrate improvements in pLDDT and ipTM (**fig. S17**). **(D)** HalluDesign for unconditional structure generation. Protenix-based HalluDesign generated more compact structures (evaluated by protein radius of gyration, Rg) than the AF3-based HalluDesign. **(E)** Incorporating AF3 for Protenix-based Halludesign is essential for producing high-quality monomers from random sequence input. ESMFold computational metrics were used for evaluation. **(F)** HalluDesign for binder design from scratch. De novo generation of binders targeting various biomolecules, including proteins (PDL1 and FGFR2), single-stranded RNA (BIV TAR RNA), and small molecules (in silico small molecule set, **fig. S8**). Averaged computational metrics (evaluated by AF3) across multiple targets for each category show consistent improvement with each HalluDesign iteration (see **fig. S22 and fig. S23**).

Despite the absence of initial protein–ligand interactions and presence of ligand-protein steric clashes inherent to the starting placement, HalluDesign simultaneously co-optimized the placement and configuration of the ligand, as well as the sequence and structure of the protein. This iterative refinement consistently converged to high-confidence complexes with high in-silico success rates (**fig. S13**). This success rate was inversely proportional to ligand size and polarity: for small, hydrophobic ligands, the rate exceeded 60% (**fig. S14A**), while it was considerably lower for larger, polar targets. This direct co-optimization approach bypasses the need for initial docking steps commonly used with previous methods (*39*). Furthermore, HalluDesign achieved significantly higher in silico success rates on these tasks compared to both diffusion-based models like RFdiffusion all-atom (*36*) and backpropagation methods like BoltzDesign1 (*21*) (**fig. S14A**). We also investigated alternative initialization strategies, such as pocket centre initialization and direct hallucination initialization, which exhibited varying declines in success rates across most ligand targets (**fig. S14B,C**).

Antibody and nanobody frameworks serve as ideal predefined scaffold platforms for targeting a wide range of molecule, as they provide highly conserved scaffolds that presents their hypervariable loops (CDRs) (**Fig. 3B**). De novo design antibody and nanobody remains challenging due to the flexible loop-dominated interfaces, and we hypothesized that HalluDesign could leverage AlphaFold3’s strong performance on antibody–protein complexes prediction to facilitate antibody and nanobody design (*13, 16*). Using Herceptin as a representative scFv framework and h-NbBcll10FGLA as a representative nanobody framework (*13*), HalluDesign successfully design and optimized CDR loops from Gaussian noise and polyglycine sequence input (**Fig. 3B** and **fig. S15A,B**). For protein specific antibody design, the target protein and the antibody framework were provided as independent templates. By randomly initializing the CDR loop coordinates and aligning their center with that of the target hotspots, HalluDesign generated antibody structures featuring CDR-specific, favourable interactions with the target (**fig. S15C**). Both structural accuracy and confidence metrics improved steadily throughout iterative optimization cycles. When targeting a phosphorylated peptide, HalluDesign produced complementary interactions between antibody CDR loops and the peptide, with the phosphate group buried in the binding interface and forming favourable charge–charge interactions (**fig. S15D**). To further allow for user-defined epitope targeting, we integrated Protenix instead of AF3 as the foundation model, leveraging its contact constraint features to guide specific epitope binding (**fig. S16**). Collectively, these results demonstrate HalluDesign’s capability for designing antibodies and nanobodies that target specific epitopes and post-translational modifications.

### HalluDesign for from scratch protein design

We next extended HalluDesign to the challenging task of de novo design from scratch. Instead of starting from folded protein, we initialized the design with purely random amino acid sequences with Gaussian noise as initial coordinates. A preliminary but plausible conformation is generated and subsequently refined through standard HalluDesign cycles. We first applied this approach to design cyclic peptide binders against protein targets (**Fig. 3C** and **fig. S17**). Two modifications are adapted to the general workflow: the target protein structure was provided as a template, and the peptide’s position index in AF3 was adjusted to enforce a head-to-tail cyclization (*40, 41*) (**fig. S17A**). By aligning the center of the random initialized peptide coordinates with that of the target hotspots, HalluDesign can generates cyclic peptides that form favorable interactions with the desired targeting site (**fig. S17B**). By leveraging AF3-style model’s ability to model covalent interactions, our approach enables the direct incorporation of unnatural amino acids (UAAs) into cyclic peptide design (*42, 43*). This enables the design of peptides capable of side-chain-to-side-chain cyclization, and covalent peptide binders that engage target cysteine residues through the incorporation of UAAs (**fig. S18**). Consequently, HalluDesign significantly expands the chemical diversity accessible for cyclic peptide binder design, achieving versatility unattainable by previous methods (*44, 45*).

However, when extending this de novo strategy to proteins substantially larger than peptides, we observed that the, the standard AlphaFold3-based HalluDesign consistently collapsed into low-complexity conformations, typically displaying as single extended helices (**Fig. 3D**). We hypothesize that this failure mode arises because random initializations lie too far outside the learned manifold of natural proteins. To address this, we replaced the AF3 engine with Protenix, an AF3-style structure prediction model augmented with protein language model (PLM) embeddings (*19*). We reasoned that the rich contextual information provided by the PLM could compensate for the lack of semantic signals in random initial sequence and effectively smooth the rugged protein sequence–structure landscape, allowing HalluDesign to find a convergence path even from highly “out-of-manifold” starting points. We observed that Protenix-based HalluDesign enabled the unconditional generation of monomeric proteins with diverse topology across a wide range of protein lengths (**fig. S19**). Notably, the generated structures exhibited substantial dissimilarity to natural proteins (see more in Supplementary Methods), highlighting the inherent de novo generative capabilities of this framework. Furthermore, by enforcing sequence identity constraints across multiple chains, HalluDesign readily produced homo-oligomers as well (**fig. S20**).

While Protenix enables the ab initio generation, we observed that its performance in subsequent refinement cycles remains inferior to that of AF3. To synergize the strengths of both foundation models, we established a ‘cross-model’ design protocol: employing Protenix for the initial generation of diverse backbone topologies, followed by iterative HalluDesign refinement cycles that alternate between AF3 and Protenix. Notably, this interleaved strategy yielded simultaneous improvements in evaluation metrics assessed by both models compared to using each single model alone (**Fig. 3E** and **fig. S21**), underscoring the robustness of the consensus optimization strategy.

Finally, we leveraged this generative capability to design binders from scratch against diverse biomolecular targets. Unlike optimization or scaffold-based approaches, this strategy operates without any initial binder structure, requiring only the target structure. To validate the multi-modal generative capabilities of our framework, we selected three distinct target classes for proof-of-concept: the protein targets, ssRNA target and small molecules (**Fig. 3F**, **fig. S22 and fig. S23**). Our cross-model HalluDesign successfully generated a diverse set of structures with favorable interactions with each target. Trajectory analysis (**Fig. 3F, fig. S22 and fig. S23**) reveals that these initial designs rapidly evolved into high-confidence binders with high structural self-consistency. The successful extension of protein binder design to a wide range of biomolecules demonstrates that HalluDesign provides a unified framework for generating binders against diverse classes of biological targets.

### Experimental validation of ligand binding proteins

To thoroughly test our HalluDesign pipeline for small molecule binder design, we selected a diverse set of ligands. This set include key endogenous small molecules characterized by multiple rotatable bonds and polar groups (**Fig. 4A** and **table S1**), including cyclic adenosine monophosphate (cAMP), adenosine triphosphate (ATP), and adrenaline (AD); small molecules with relatively large, rigid, and hydrophobic features, such as cholic acid (CA) and the HIV drug dolutegravir (DTG); as well as the amino acid tryptophan (TRP). We also extended our HalluDesign approach to metal ion ligand zinc (Zn²⁺). These targets collectively represent a broad diversity of ligand sizes and chemical properties. NTF2 scaffolds were used for ATP, CA, DTG, and AD; pseudocycles for Zn²⁺; and both scaffold types for cAMP and TRP. Protein-ligand complex structures were initialized using the “black hole initialization” strategy and HalluDesign was applied to generate the final protein-ligand complexes (**fig. S24**). Unlike conventional high-throughput approaches that require designing and screening large numbers of candidates (*38, 46*), HalluDesign enabled binder discovery in a low-throughput setting. For each small-molecule target, a maximum of 16 designed proteins were experimentally characterized, most of which were successfully expressed in *E. coli*. ITC measurements confirmed one or more binders for every target (**Fig. 4B**, **fig. S25**, and **table S2**). Mutational analysis of the ligand binding pocket residues confirmed both the binding mode and the accuracy of the designed interactions (**fig. S26A**). Cross-binding assays demonstrated high ligand specificity (**fig. S26B,C**); for example, the TRP binder did not bind to the other aromatic amino acids, such as tyrosine or phenylalanine, highlighting the platform’s potential for designing amino acid-specific binders. Similarly, the AD binder exhibited minimal binding to its functional homolog, norepinephrine (NE), which differs from AD by a single methyl group, underscoring the specificity achieved by our designs.

**Fig. 4.**
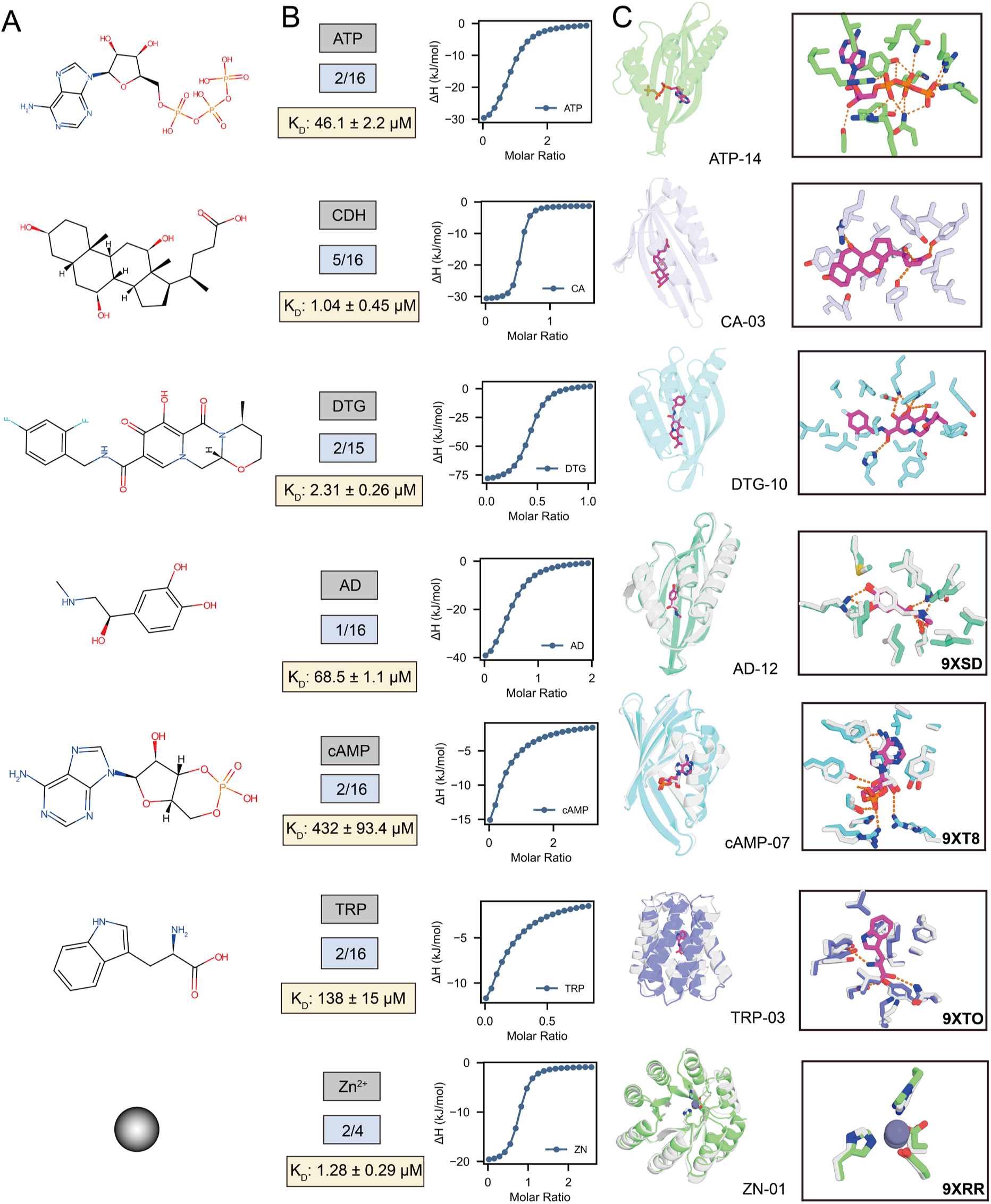
Experimental validation of ligand binders generated by HalluDesign. **(A)** HalluDesign successfully generates binders for a range of targets. **(B)** Experimental success rates, binding affinities and ITC profiles for the representative designs shown in (**C**). **(C)** Design models and crystallographic validation of HalluDesign-generated binders. Crystal structures were obtained for cAMP, AD, Zn²⁺, and TRP binders. Superposition of experimentally resolved structures (grey) with design models (colored) demonstrates high atomic accuracy for these targets; the TRP binder structure was determined in the absence of ligand. Electron densities of the ligands and their neighboring residues are shown in **fig. S27**. For designs without crystal structures, only the computational models are shown.

We successfully obtained crystal structures of protein-ligand complexes for cAMP, AD and Zn²⁺ (**Fig. 4C** and **fig. S27**). The cAMP-binder complex was resolved at 2.8 Å, with clear electron density for cAMP and the overall structure closely matching the design model (Cα RMSD of 0.79 Å for the protein and 0.98 Å for cAMP using the protein as reference). The sidechain configurations of pocket residues were accurately recapitulated. Notably, all seven designed hydrogen bonds between the protein and cAMP were well observed in the crystal structure. The AD binder complex was resolved at 2.2 Å resolution, showing atomic-level agreement with the design model (Cα RMSD of 0.63 Å for the protein and 0.32 Å for AD). In both the design model and crystal structure, the binder interacts with AD through several hydrogen bonds. The terminal methyl group of AD is tightly packed with surrounding hydrophobic residues, which underlies the high specificity for AD over NE. The Zn²⁺ binder complex was resolved at 1.6 Å, also demonstrating overall agreement with the design model (Cα RMSD of 0.81 Å for the protein and 0.75 Å for zinc). Zinc adopts a tetrahedral coordination geometry in both the design model and the crystal structure, with all four coordinating residues precisely positioned. We also obtained an apo structure for the TRP binder at 1.9 Å resolution, which also closely match with the design model (Cα RMSD of 0.79 Å). Pocket residue sidechain orientations in this apo structure mirrored those in the design model. The high experimental success rate, combined with point mutation analysis of the binding pocket and the atomic agreement between design models and crystal structures, showcase the efficiency and accuracy of HalluDesign in creating specific protein-ligand interactions.

### Experimental validation of designed protein-protein interactions

We applied HalluDesign to rescue previously unsuccessful designs in the RFdiffusion protein–protein interaction (PPI) design task (*35*). For these experiments, we truncated the AlphaFold3 diffusion trajectory and applied structure conditioning during the structure update stage to better preserve similarity between the final complex and the initial design model. The resulting design models exhibited improved computational metrics and exhibited lower RMSD deviation from the original input structures (**Fig. 2E** and **fig. S7**). For experimental validation, we selected 16 rescued designs each for PD-L1 and IL-7Rα, with each candidate derived from a previously failed design (**Fig. 5A**). Biolayer interferometry (BLI) assays demonstrated that 15 out of 16 PD-L1 binders successfully bound their target, while 8 out of 16 IL-7Rα binders showed binding to IL-7Rα (**fig. S28**). Overall, most of these designs achieved low nanomolar affinities for their respective targets. Interface disruption mutations totally abolished the binding signal, and these results validated the design configurations (**fig. S29**). The high success rate of these rescue experiments underscores the efficiency of the HalluDesign approach in optimizing protein–protein interactions.

**Fig. 5.**
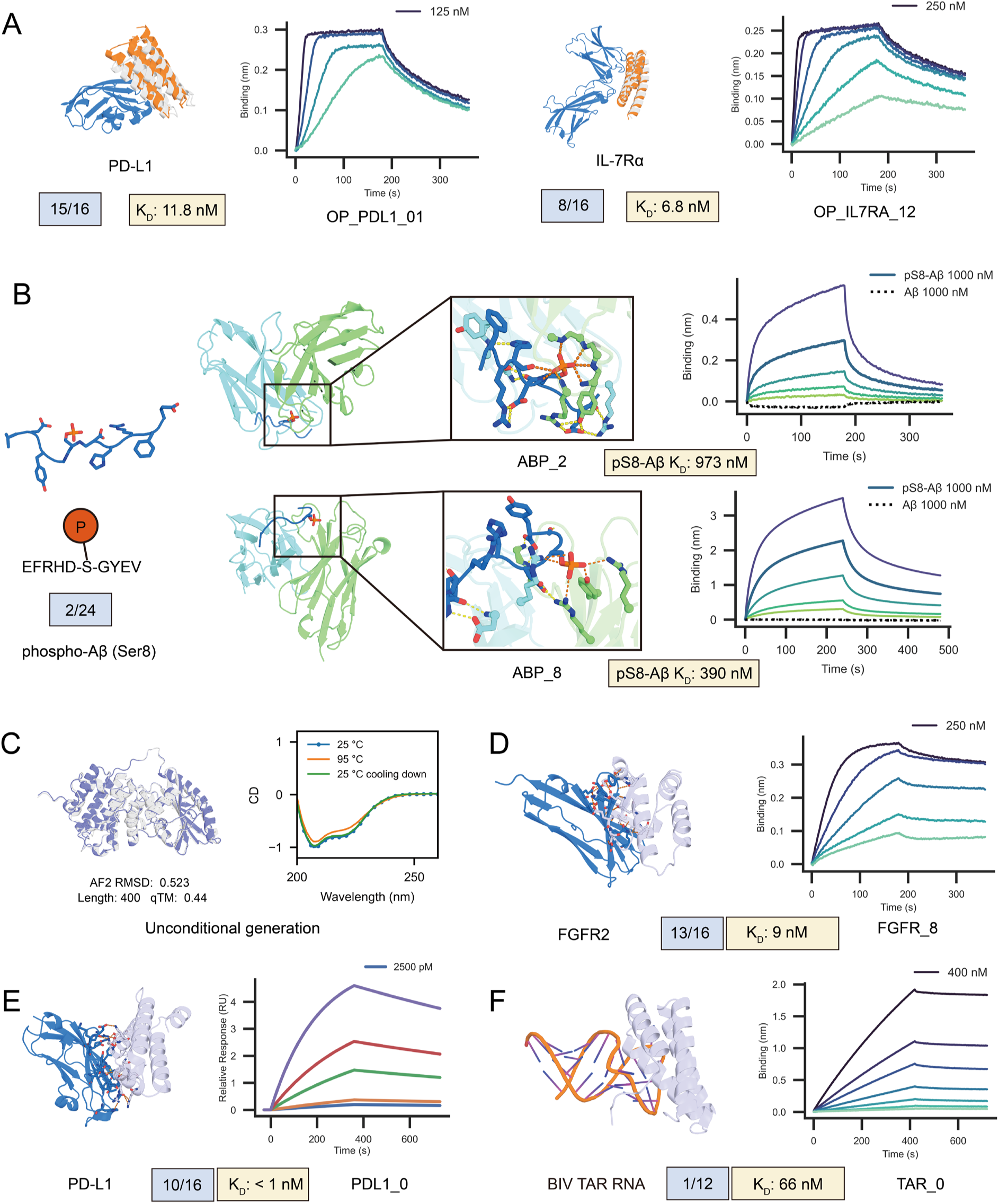
Experimental validation of protein interface optimization, antibody design, and de novo protein generation using HalluDesign. **(A)** Rescue of failed RFdiffusion binders by HalluDesign. HalluDesign efficiently optimizes non-binding scaffolds, generating experimentally validated binders for PD-L1 and IL-7Rα targets with high success rate. White denotes the original scaffold, blue the target protein, and orange the optimized binder. **(B)** HalluDesign for phosphorylation-specific antibody design. Designed scFvs (green/blue) target Aβ peptide with the phosphorylated serine. BLI assays demonstrated specific binding to the phosphorylated Aβ, whereas no binding signal was detected for the native peptide. **(C)** Experimental characterization of a representative de novo protein monomer generated by HalluDesign. The final design model shown in color, while the predicted structure from ESMFold is grey. CD analysis demonstrated high thermal stability. Additional experimental results for other monomers across various lengths are provided in **fig. S30**. (**D, E**) HalluDesign for de novo protein binder design. Experimental validation for two distinct targets, PD-L1 and FGFR2. The HalluDesign-generated PD-L1 binder exhibited high binding affinity as measured by surface plasmon resonance (SPR). HalluDesign demonstrated high design success rates; BLI results for additional successful designs are shown in (**fig. S31)**. (**F**) HalluDesign for RNA binder design. BLI results for the HalluDesign generated binder targeting BIV TAR RNA. For the BLI or SPR titrating experiments, only the highest concentration is labeled; the remaining concentrations follow a two-fold serial dilution.

Targeting amyloid β (Aβ) with high specificity remains a crucial strategy in Alzheimer’s disease (AD) therapy (*47*). Phosphorylation at Ser8 (pS8-Aβ) promotes toxic oligomer formation and accelerates amyloid pathology. Designing antibodies that selectively recognize this phosphorylated form could enable targeted intervention against the neurotoxic Aβ species (*48*). Starting from the Herceptin scFv framework, HalluDesign successfully generated new CDR loops that form favourable interactions with phosphorylated Aβ peptide. Unlike conventional methods that rely on predefined target peptide positions or second structure (*14, 49*), HalluDesign allows for free sampling of peptide conformation during the antibody design process. 24 scFv designs were selected for experimental characterization and 16 were found to be readily expressed in *E. coli*. Experimental results demonstrated that two of the designed scFvs could specifically bind to the phosphorylated form of Aβ, with no detectable interaction with native Aβ (**Fig. 5B**). These findings indicate that the phosphate group is critical for antibody-peptide interaction, aligning with our design model where the phosphate group is deeply buried within the interface and engages in electrostatic interactions with adjacent residues on the antibody. The successful design of highly specific, phosphorylated-peptide-targeting antibodies underscores HalluDesign’s potential for developing antibodies against post-translational modifications (PTMs).

### Experimental validation of from scratch generated proteins

To evaluate the HalluDesign’s capability for from scratch protein generation, we experimentally validated 14 monomeric proteins ranging in length from 70 to 400 residues. All but one were successfully expressed in *E. coli* with high expression yield. Size-exclusion chromatography confirmed their monomeric elution, and circular dichroism assays demonstrated their exceptional thermostability (**Fig. 5C** and **fig. S30**). We next employed HalluDesign to the de novo design of protein binders with biomedical relevance—specifically targeting PD-L1 and FGFR2, which are key proteins in immunotherapy and cancer treatment (*50, 51*). For each target, we experimentally tested 16 de novo designed binders and observed high success rates: 10 out of 16 PD-L1 binders showed binding signals in BLI experiments, with the strongest affinity reaching picomolar binding in low concentration surface plasmon resonance (SPR) experiment, and 13 out of 16 FGFR2 binders showed binding, with the best achieving digital nanomolar affinity (**Fig. 5D,E** and **fig. S31**). Interface mutation experiments further confirmed that the binding modes were consistent with the design models (**fig. S32**). Extending HalluDesign to nucleic-acid targets, we designed protein binders for TAR (the trans-activation response element) RNA, a critical regulatory element in bovine immunodeficiency virus (BIV) transcription (*52*). Designing specific binders against TAR RNA holds potential for the inhibition of viral transcription. Among the 12 candidates selected for experimental validation, we identified one binder with a measured affinity of 66 nM in BLI experiments (**Fig. 5F**). These results demonstrated that our framework naturally extended beyond proteins and could handle nucleic-acid targets as well. Collectively, these robust experimental success rates and high binding affinities highlight the power of HalluDesign as a platform for generating effective binders across a broad range of biomolecular targets.

## Discussion

In conclusion, we present HalluDesign, a versatile all-atom framework that fundamentally reframes the utility of state-of-the-art structure prediction models for protein design. Rather than employing AF3-style models as passive, external ‘oracles’ to filter candidate designs, HalluDesign harnesses the hallucination effect intrinsic to these models to enable iterative sequence–structure optimization. By smoothing the rugged sequence–structure landscape, our approach facilitates efficient co-optimization, resulting in rapid convergence without the need for costly backpropagation or model fine-tuning. Leveraging the versatile biomolecular modeling capabilities of AF3-style models, HalluDesign enables the optimization and design of diverse biomolecular structures and interactions. Looking forward, the framework is designed to be extensible to alternative AF3-style models, such as Boltz-1 and Chai-1, together with ongoing advances in structure prediction, the capabilities of HalluDesign will be further enhanced.

Our structure conditioning strategy provides a straightforward way to control the sequence–structure co-optimization process, enabling HalluDesign to be employed across a wide range of scenarios—from refining existing structures to fully de novo protein generation from random sequences. In optimization tasks, HalluDesign consistently enhances computational metrics for structures generated by other state-of-the-art methods. Notably, experimental characterization demonstrates that HalluDesign can successfully rescue previously failed binders generated by RFdiffusion with a high success rate. However, for candidates that already exhibit detectable binding, further experimental evaluation is necessary to determine whether our improvements in computational metrics translate into significant affinity gains. Additionally, more precise control over structure conditioning, such as focusing on motif geometry or the configuration of catalytic functional groups, will extend HalluDesign’s applicability to tasks like motif scaffolding and de novo enzyme design.

The HalluDesign framework is generalizable beyond protein design and, in principle, can be adapted to design any biomolecule modeled by AF3-style predictors. For instance, substituting protein sequence design models with nucleic acid sequence design models would enable the design of new DNA and RNA structures (*53*), while integrating small molecule engineering methods could facilitate novel small molecule drug discovery (*54, 55*). Ultimately, these advances will empower de novo design beyond proteins—encompassing nucleic acids, small molecules, and complex biomolecular assemblies—and pave the way for unprecedented innovations in synthetic biology, biotechnology, and therapeutic discovery.

## Acknowledgments

We appreciate the members of Cao’s labs for their valuable discussions. We extend our special thanks to Dang Lab in Westlake University for valuable discussion about covalent binding cyclic peptide. This work was supported by State Key Laboratory of Gene Expression. We thank the Protein Characterization and Crystallography Facility of Westlake University for help in sample analysis; the Mass Spectrometry & Metabolomics Core Facility of Westlake University for sample analysis; the Westlake University HPC Center for computation assistance.

## Author contributions

Conceptualization: CW, MF, LC

Methodology: MF, CW, LC

Investigation (computational): MF, CW, PC

Investigation (experimental): JS, FL, QJ, ZW, ZC, YW, MF, QH

Visualization: MF, ZZ, YZ, YK

Funding acquisition: LC

Supervision: LC

Writing – original draft: MF, CW, LC

Writing – review & editing: LC, CW, MF

## Data availability

We have obtained four PDB IDs for these structures: AD_12 (00009XSD), ZN_01 (00009XRR), cAMP_07 (00009XT8), and TRP_03 (00009XTO).

## Code availability

All source code developed in this study are available at:

CoDP (https://github.com/MinchaoFang/CoDP) and

HalluDesign (https://github.com/MinchaoFang/HalluDesign).

## Supplementary Materials

Materials and Methods

Figs. S1 to S33

Tables S1 to S5

---

**Fig. S1.**
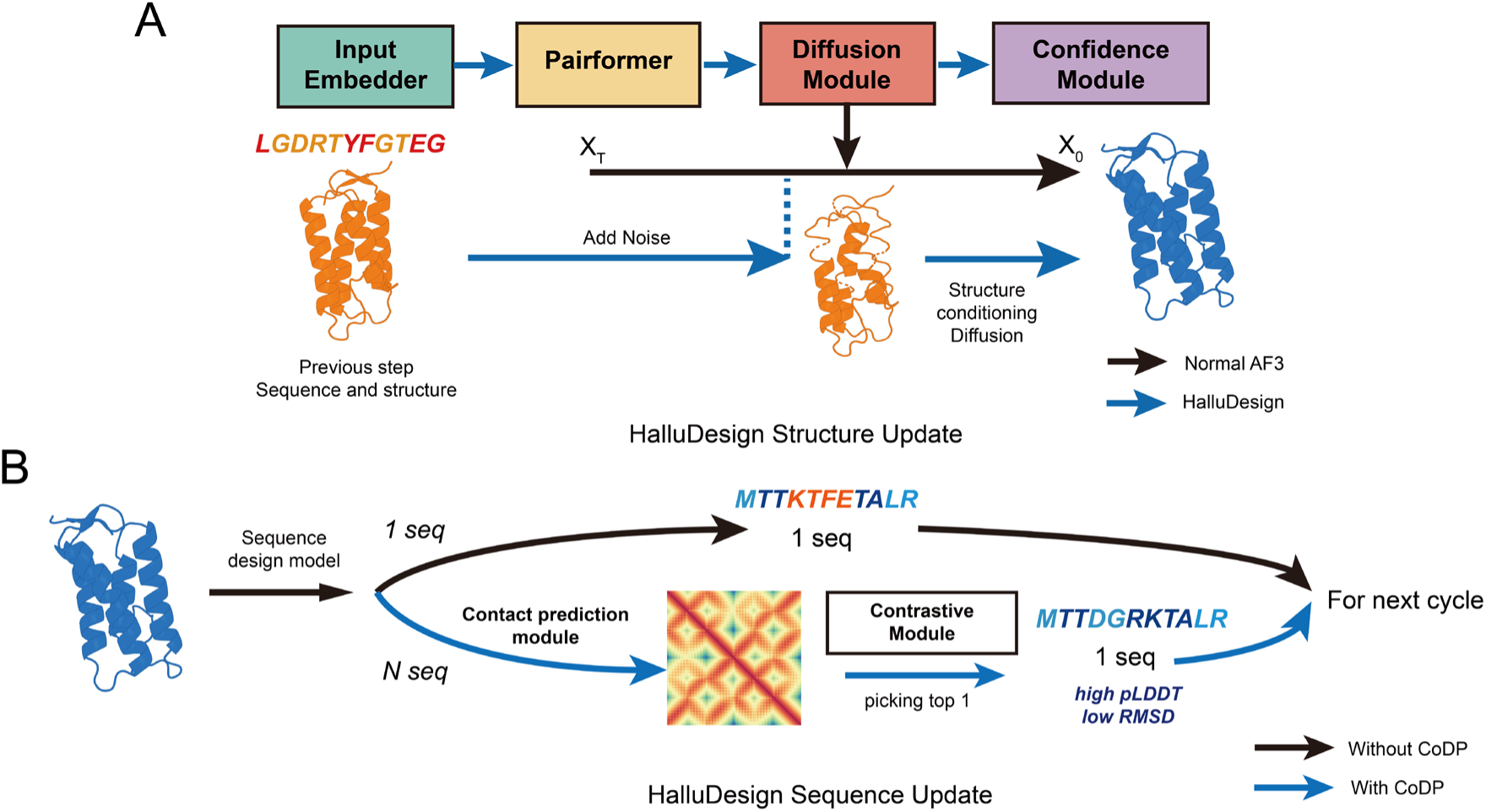
Overview of the structure and sequence update stages of HalluDesign. (**A**) Truncated diffusion approach within the AlphaFold3-style model. After sequence inputs are processed through the Input Embedder and Pairformer, a user defined noise level is applied to the structure from the previous step. This noised structure (X_T_) serves as the input for the reverse diffusion process in the Diffusion Module. The module then generates an updated structure (X_0_), which is subsequently passed to the Confidence Module for confidence evaluation. This structure conditioning allows precise control over sampling in structure space. (**B**) Sequence update stage. A sequence design model generates a new sequence for the updated structure. Optionally, multiple candidate sequences can be generated and ranked using the CoDP model, which selects the optimal sequence based on contact prediction and contrastive learning, resulting in improved sequence confidence and consistency.

**Fig. S2.**
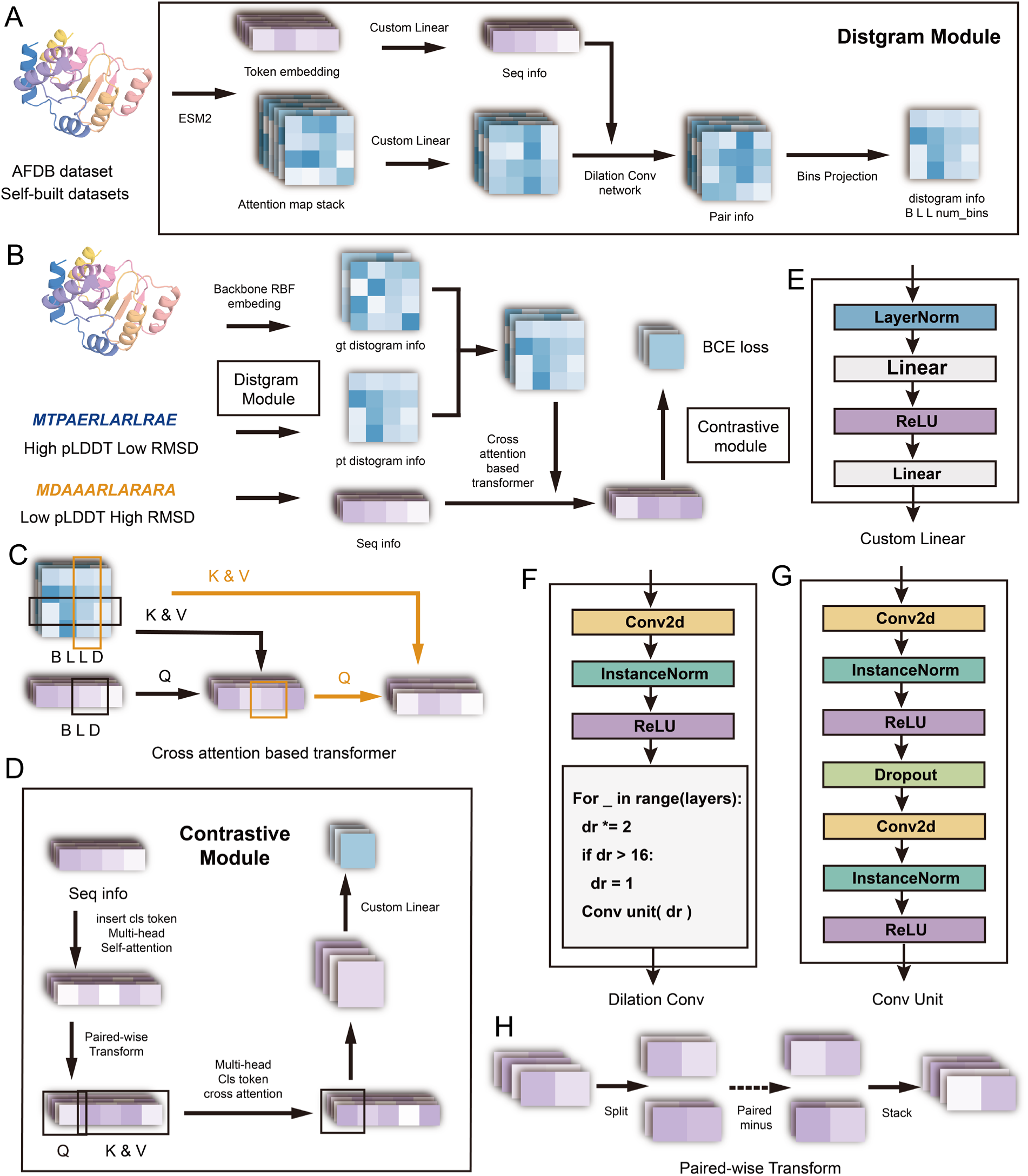
Detailed architecture of the CoDP model. The CoDP framework consists of a distogram prediction module and a contrastive scoring module, using ESM2 embeddings for feature extraction. (**A**) Distogram prediction module: Predicts inter-residue distances using a dilated convolutional network with 8 distance bins for efficient and high-resolution modeling. (**B**) Contrastive learning workflow: Integrates predicted and ground-truth distograms with sequence embeddings to evaluate design confidence. (**C**) Cross-attention mechanism: Fuses pairwise structural information into single-sequence embeddings for global feature aggregation. (**D**) Contrastive Module: Combines embeddings to score prediction confidence. Detailed structural components: (**D**) Contrastive Module: Combines embeddings to score prediction confidence. (**E**) Custom Linear Layer: Transforms and reduces feature dimensions for downstream processing. (**F**) Dilation Convolution Module: Expands the receptive field through cyclic dilated convolutions. (**G**) Convolutional Unit: Basic block of Conv2d, InstanceNorm, ReLU, and Dropout layers. (**H**) Pairwise-wise Transform mechanism: Captures pairwise differences to support alignment-aware scoring.

**Fig. S3.**
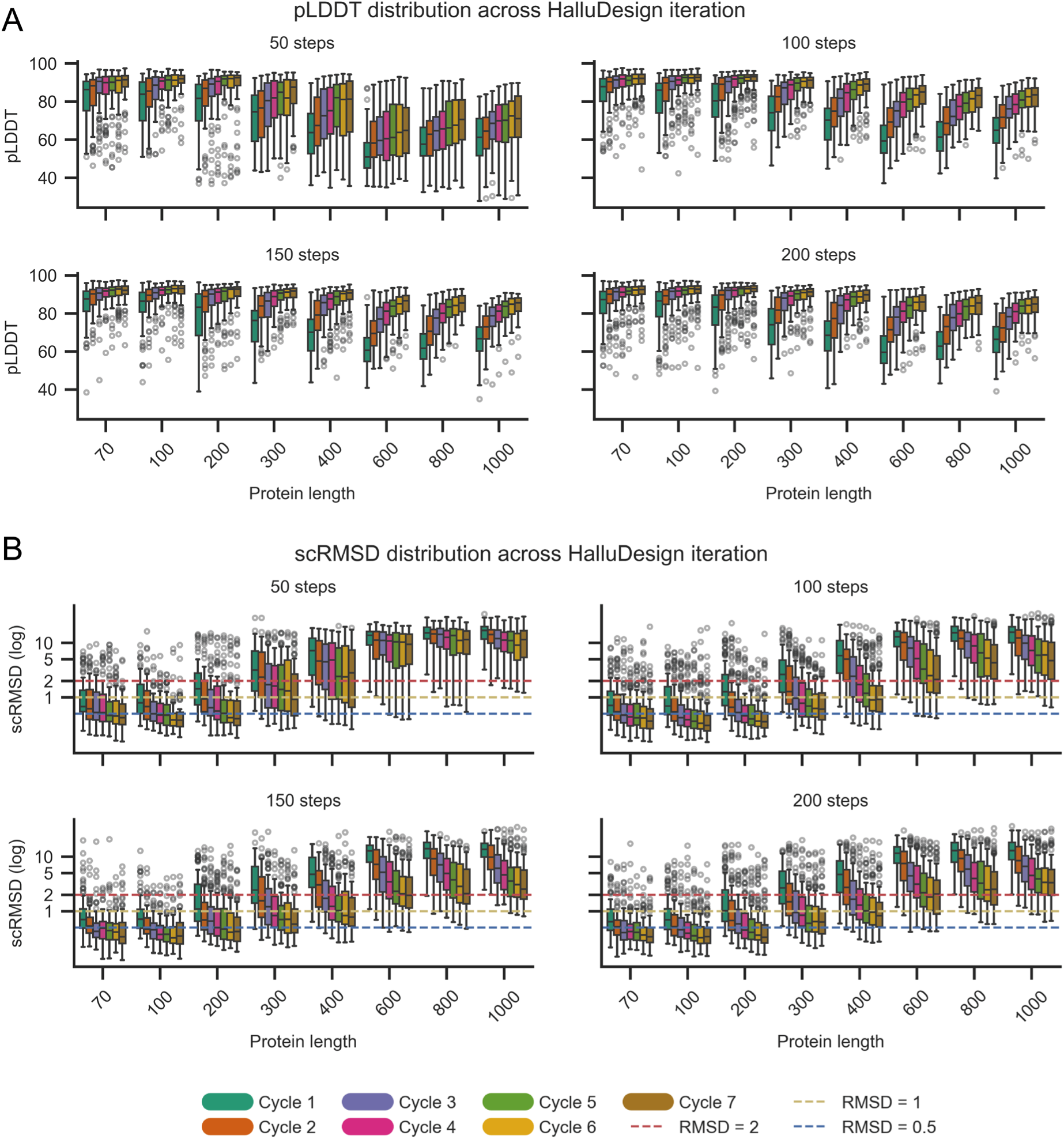
HalluDesign optimization for Chroma backbones. **(A)** pLDDT and **(B)** self-consistency Cα-RMSD distributions are shown across HalluDesign optimization cycles, indicating consistent improvement in scaffold quality. Truncating the diffusion trajectory at varying lengths (50, 100, 150 and 200 steps) results in different optimization effect, depending on the scaffold size. The default maximum length for the AlphaFold3 diffusion module is 200 steps. scRMSD and pLDDT are computed using the orthogonal model ESMFold with standard evaluation methods.

**Fig. S4.**
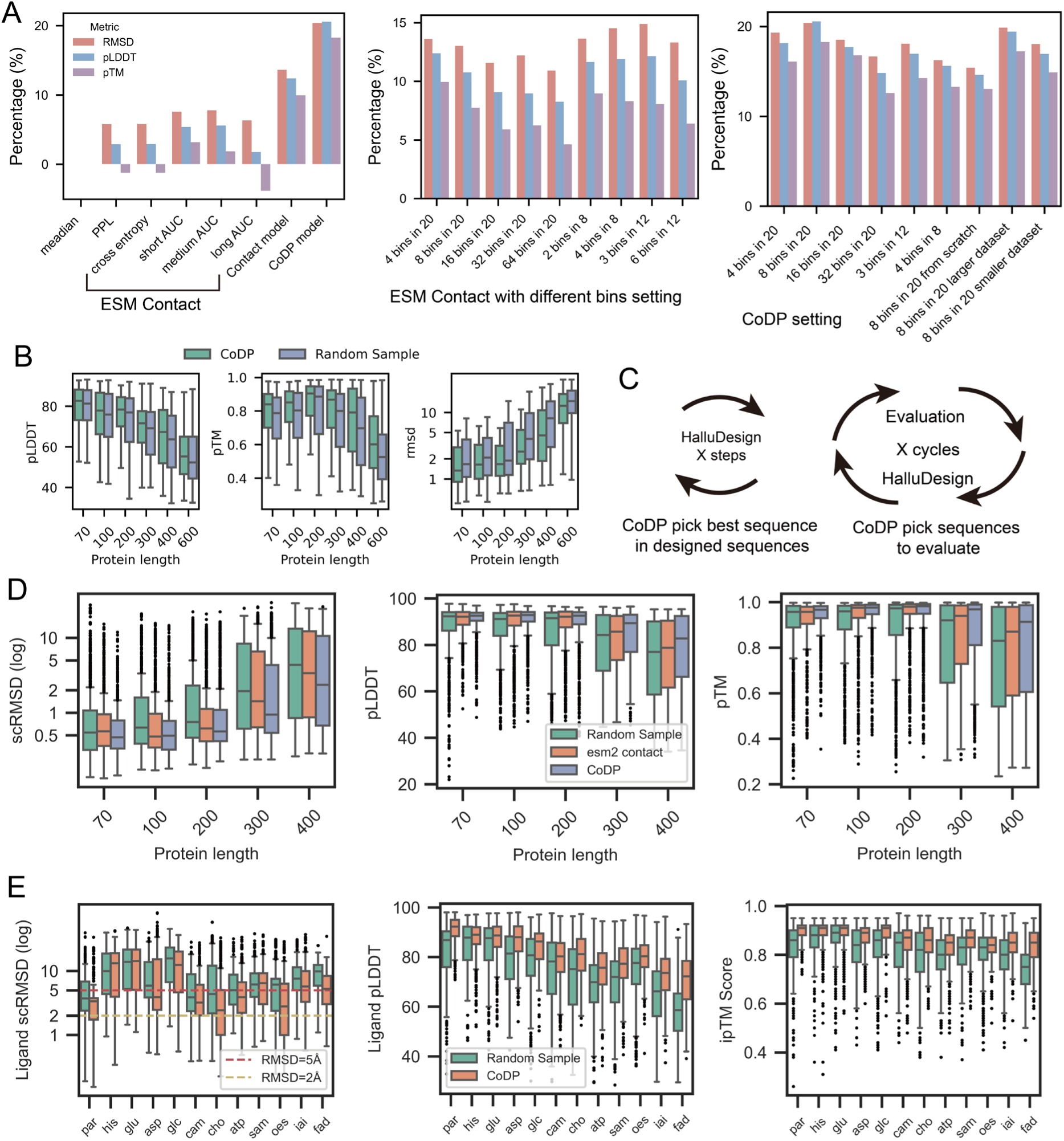
CoDP model for sequence ranking to promote optimization. **(A)** Quantitative evaluation of sequence selection strategies in our test dataset (see Methods for percentage definitions). The left panel compares the selection success rate of raw ESM-2 metrics, a trained contact model, and the full CoDP model. The middle and right panels analyze the impact of distogram binning granularity on selection accuracy, optimized separately for the standalone contact model (middle) and within the full CoDP framework (right). **(B**) Sequences selection performance. CoDP consistently identifies sequences with higher confidence scores (pLDDT, pTM) and lower RMSD compared to random selection across various protein lengths in our test dataset curated from Chroma. **(C)** Workflow showing CoDP as a sequence filter in the iterative optimization cycle and in evaluation. **(D)** Optimization performance on monomers. The CoDP strategy (purple) yields better structural metrics compared to Random Sampling (green) and ESM2 contact-based selection (orange). Designs are generated using Protenix-based HalluDesign and predicted by ESMFold self-consistency strategy with 8 sequences. **(E)** Optimization performance on ligand-binding proteins from RFdiffusion-AA with 7 cycles optimization. Box plots show distributions of Ligand RMSD, pLDDT, and ipTM, demonstrating that CoDP (orange) improves upon random sampling (green) in complex generation. Designs are generated using Protenix-based HalluDesign and predicted by Alphafold3 self-consistency strategy with 8 sequences.

**Fig. S5.**
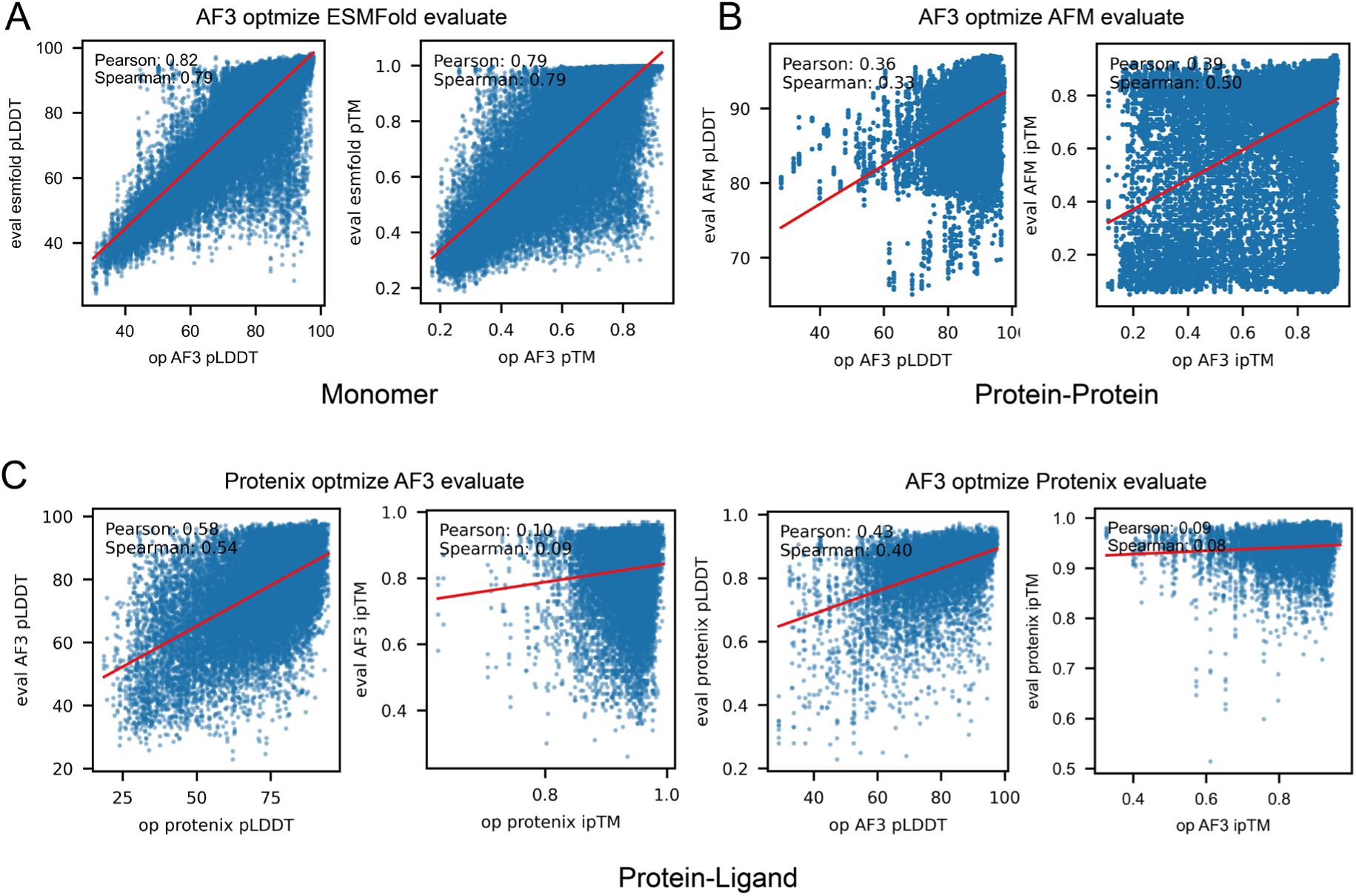
Correlation between internal confidence metric from HalluDesign optimizer and external metrics from orthogonal structure prediction models. (**A**) Monomer optimization. Scatter plots dispaly the correlation between the AF3-based HalluDesign internal confidence metrics (pLDDT and pTM at the all optimization cycle, X-axis) and the ESMFold validation metrics (predicted from the all designed sequence, Y-axis). (**B**) Protein binder optimization. Scatter plots show the correlation between the AF3-based HalluDesign internal confidence metrics (binder pLDDT and ipTM at the all optimization cycle, X-axis) and the AlphaFold-Multimer validation metrics (Y-axis). (**C**) Small molecule binder optimization. Cross-validation of confidence metrics (binder pLDDT and ipTM) using AlphaFold 3 and Protenix. One model serves as the optimizer and the other functions as the validation model. The internal confidence metrics are plotted on the X-axis and the corresponding validation metrics on the Y-axis.

**Fig. S6.**
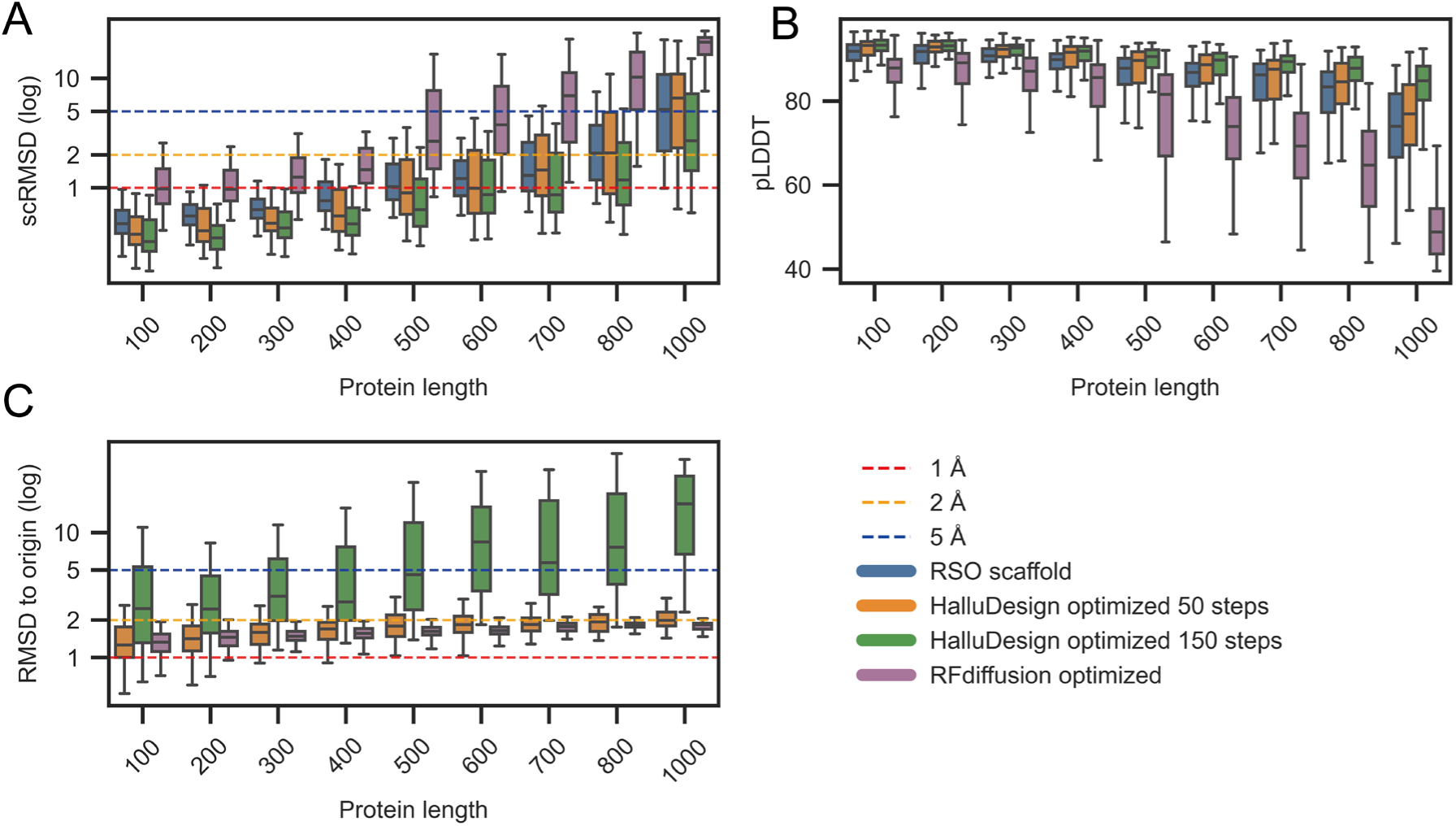
HalluDesign optimization of RSO backbones. Self-consistency Cα-RMSD (**A**), pLDDT (**B**) and Cα-RMSD relative to the original structure (**C**) are shown for various structure optimization settings and different RSO backbone lengths. For RFdiffusion, partial diffusion was performed with T = 12 with a total trajectory length of 50 steps. HalluDesign optimization was conducted over seven cycles using truncated diffusion trajectories of 50 and 150 steps, respectively. Structural evaluation was performed using ESMFold.

**Fig. S7.**
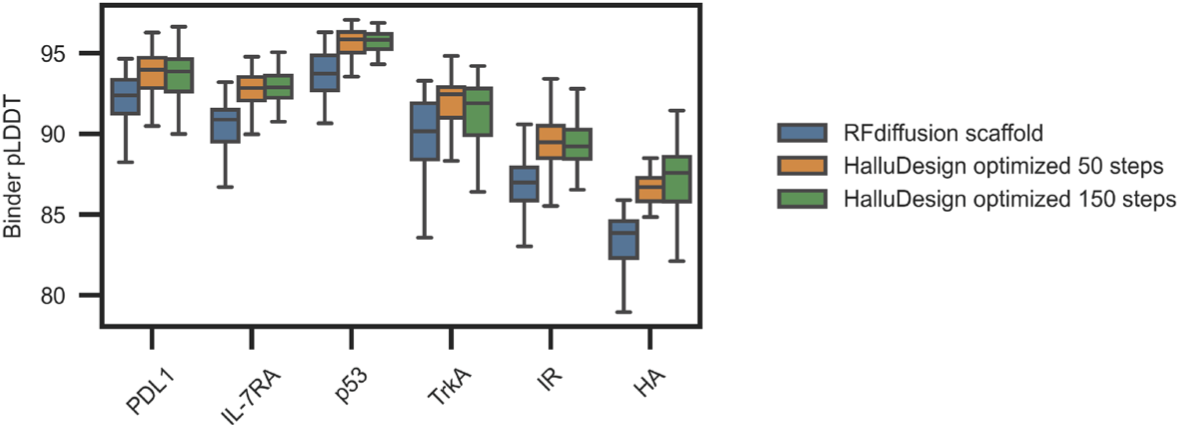
HalluDesign optimization of RFDiffusion-generated protein binders. Original binder–target complex models were from the supplementary materials of the RFdiffusion paper. Binder pLDDT distributions were evaluated using the AlphaFold2-Multimer self-consistency strategy. AF3-based HalluDesign optimization was performed over seven cycles with truncated diffusion trajectories of 50 and 150 steps, respectively. Additional evaluation metrics, including self-consistency RMSD, ipTM, and RMSD to the initial structure, are reported in Fig. 2E.

**Fig. S8.**
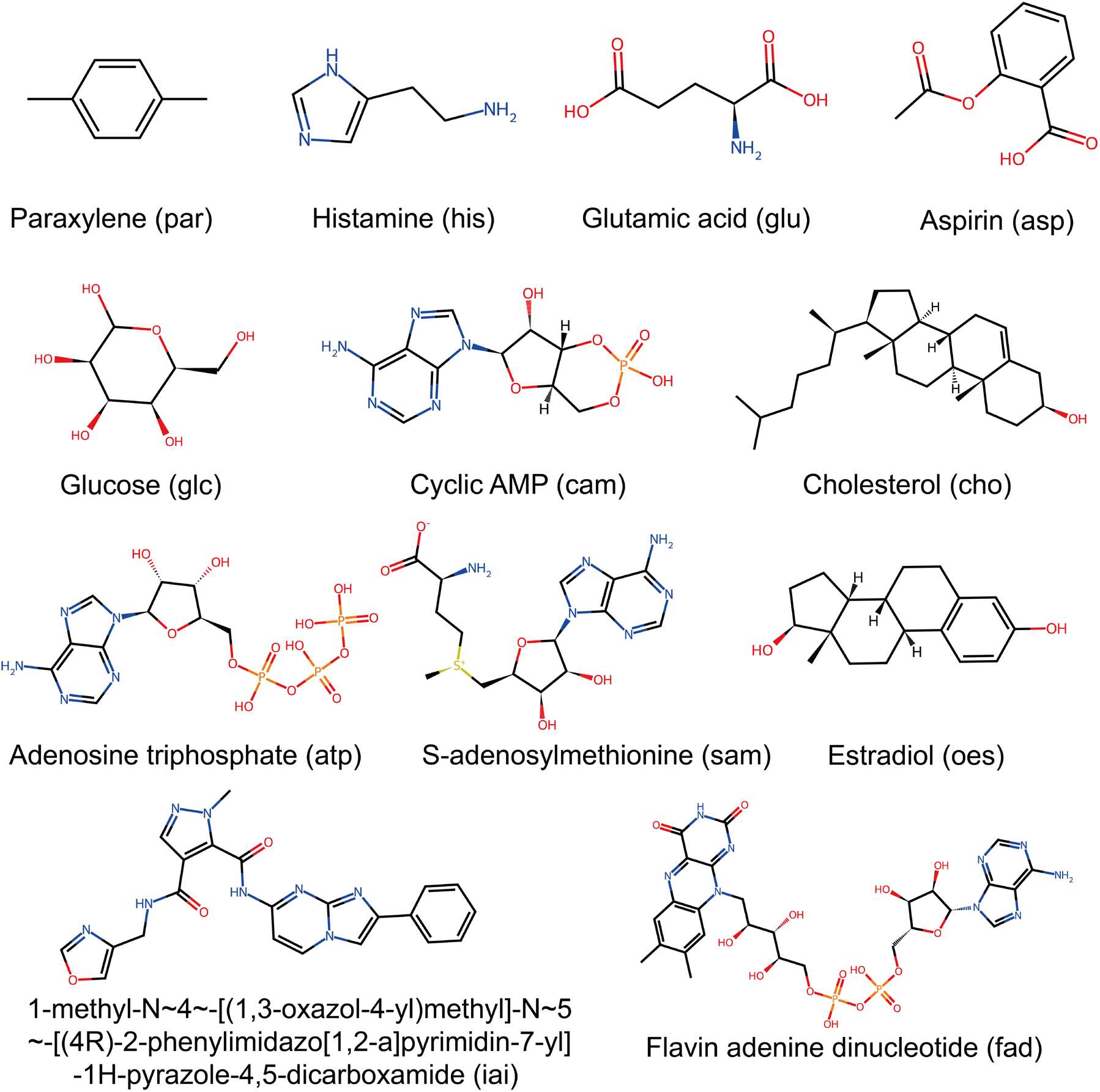
Ligand chemical structures, common names and abbreviations in the in sillico small molecule benchmark.

**Fig. S9.**
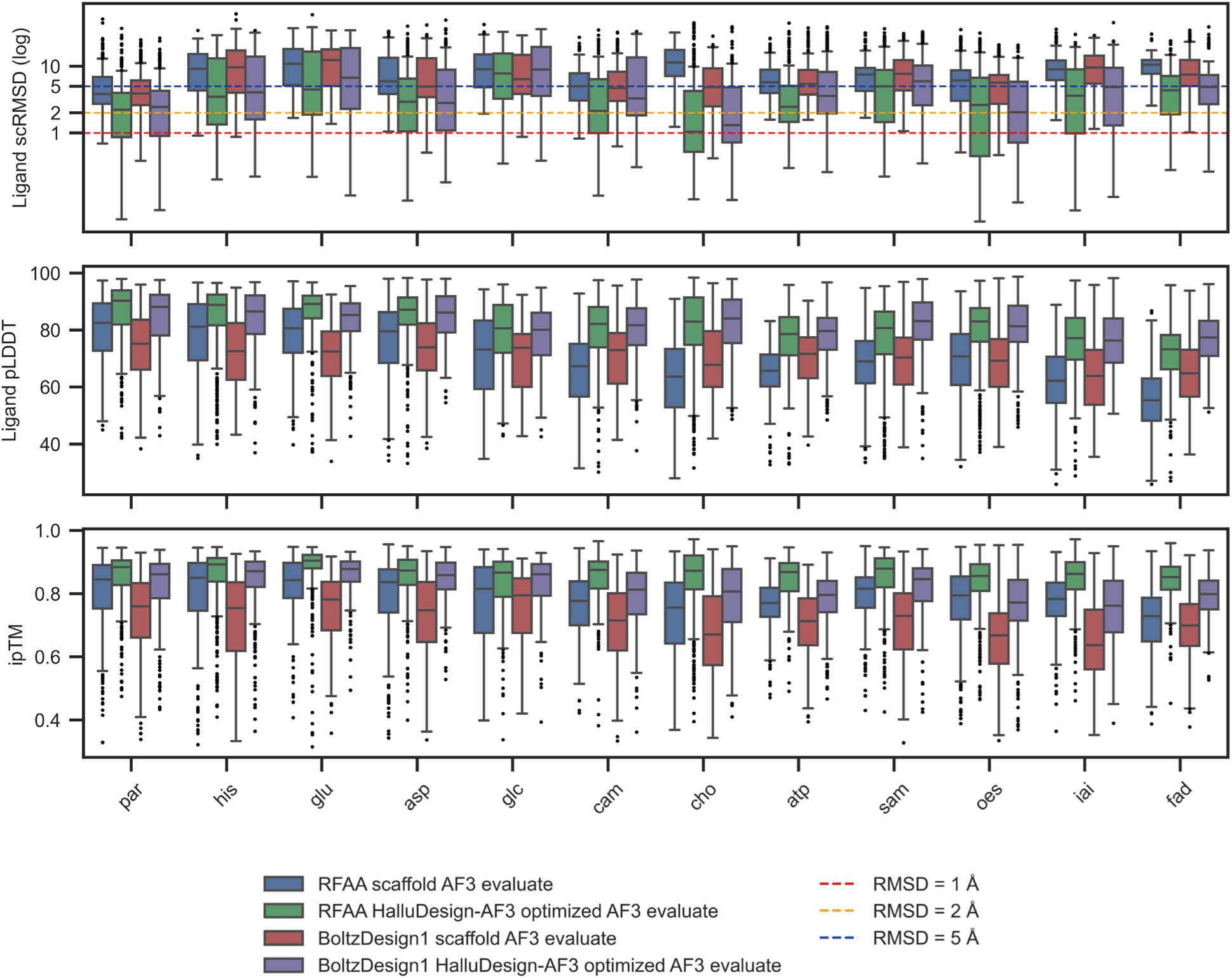
AF3-based HalluDesign optimization of small molecule binders generate by RFdiffusionAA and BoltzDeisgn1. Ligand self-consistency RMSD, pLDDT, and ipTM distributions are shown across the ligand set presented in **fig. S8**, for binders generated by RFdiffusionAA and BoltzDesign1, both before and after HalluDesign optimization. Optimization with HalluDesign was performed over seven cycles using truncated diffusion trajectories of 150 steps. Computational metrics were evaluated using the AlphaFold3 self-consistency method.

**Fig. S10.**
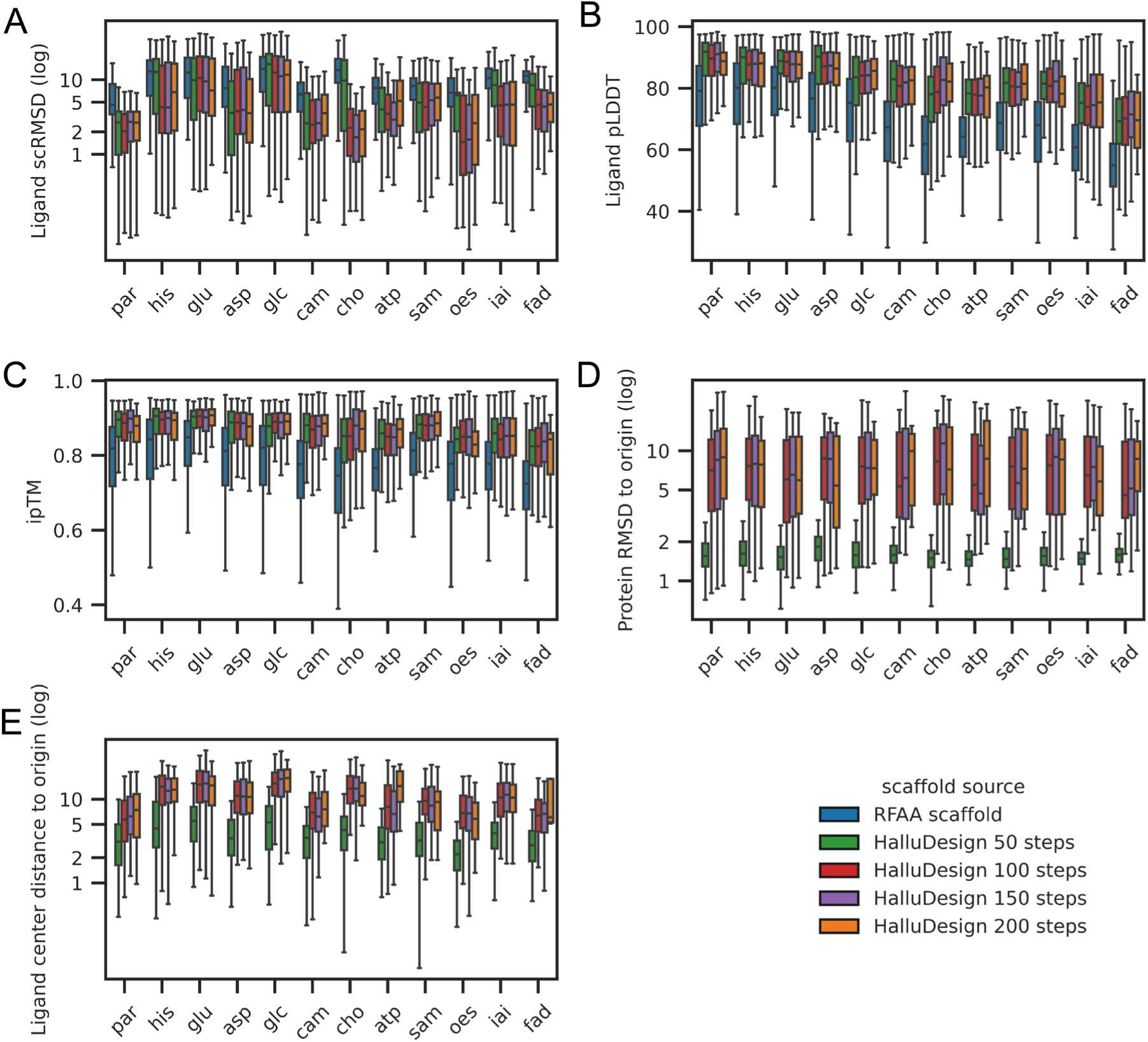
HalluDesign optimization of RFdiffusionAA generated small molecule binders using varying truncated diffusion trajectories. Ligand pLDDT (**A**), ligand self-consistency RMSD (**B**), ipTM (**C**), protein binder Cα-RMSD relative to the initial structure (**D**) and ligand center-of-mass displacement distance with respect to the protein (**E**) are shown for each small molecule target, comparing different truncated diffusion trajectory lengths (50, 100, 150, and 200 steps, with 200 steps as the default). Seven optimization cycles were performed. All metrics were evaluated using the standard AF3 self-consistency method.

**Fig. S11.**
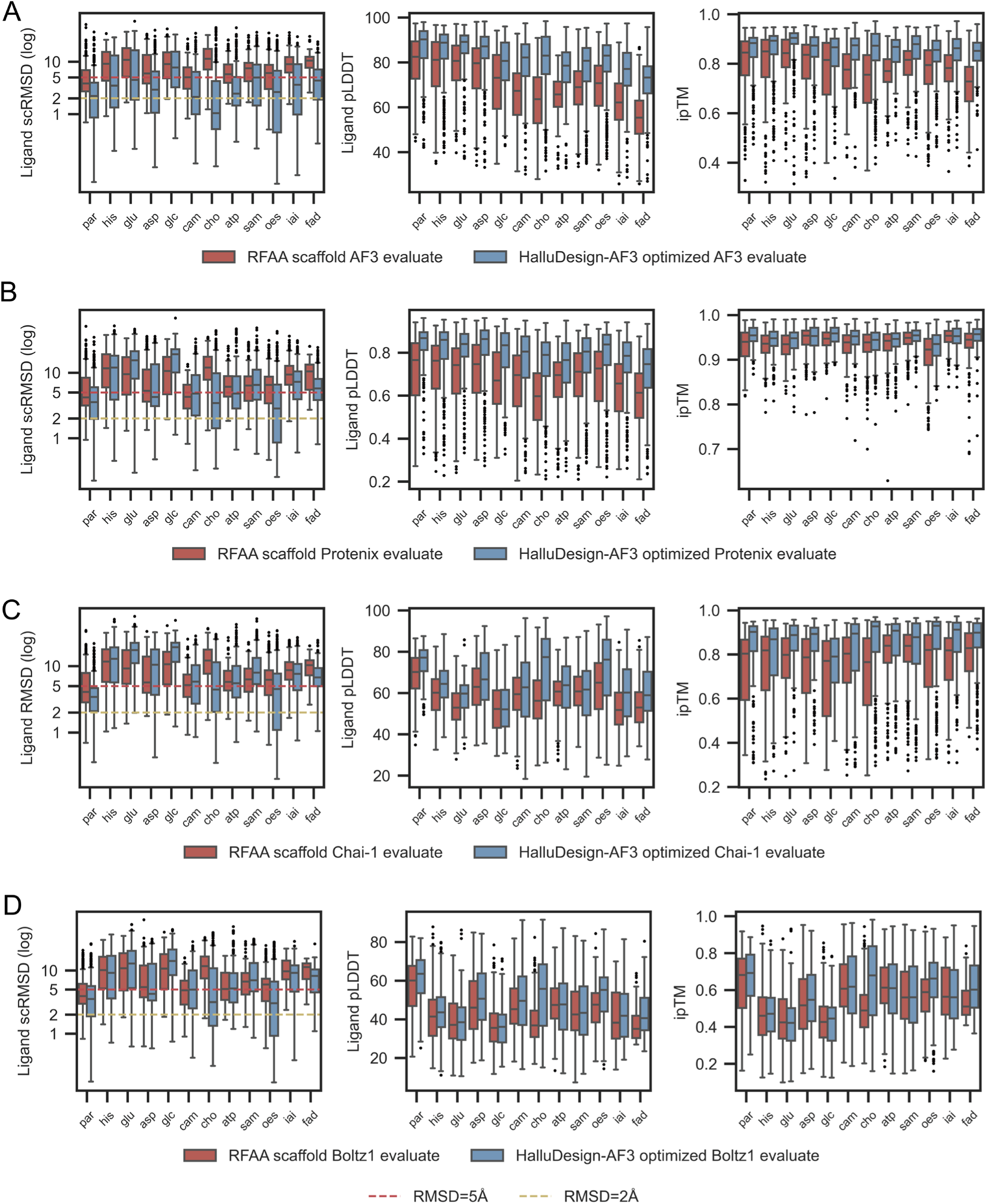
Benchmark of HalluDesign optimization on RFdiffusionAA generated small molecule binders using different stucture prediction models (AF3, Protenix, Chai-1, and Boltz-1). Ligand self-consistency RMSD, pLDDT, and ipTM distributions for RFdiffusionAA generated small molecule binders, evaluated by AF3 (**A**), Protenix (**B**), Chai-1 (**C**) and Boltz-1 (**D**), are shown before and after AF3-based HalluDesign. Seven optimization cycles of 150 diffusion steps were conducted.

**Fig. S12.**
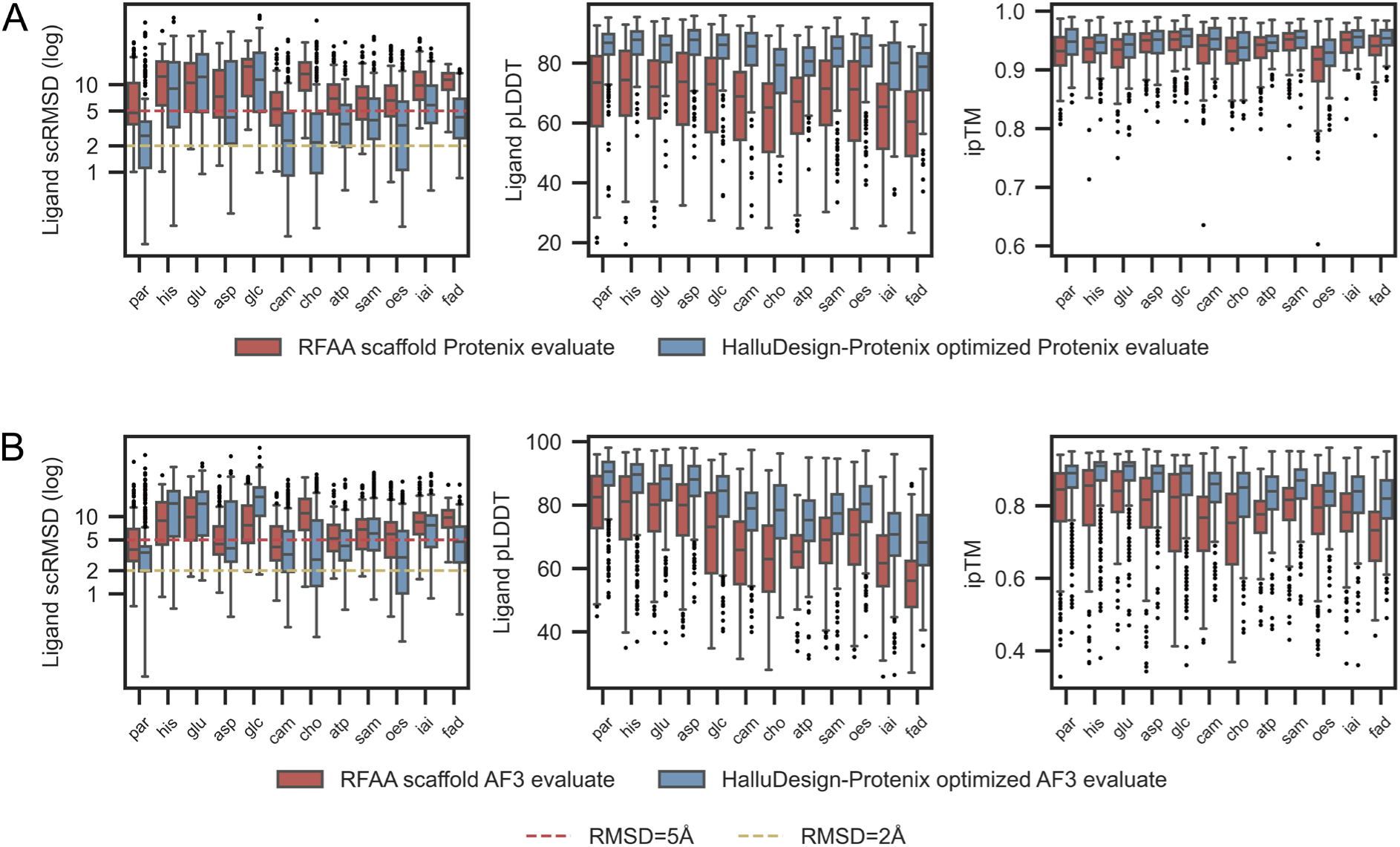
Protenix-based HalluDesign optimization on RFdiffusionAA generated small molecule binders. Ligand self-consistency RMSD, pLDDT, and ipTM distributions for RFdiffusionAA generated small molecule binders are shown before and after Protenix-based HalluDesign optimization. Seven optimization cycles were conducted, with truncated diffusion trajectories of 150 steps out of 200 total steps. Performance was evaluated using both the Protenix **(A)** and AF3 (**B**) self-consistency assessment methods.

**Fig. S13.**
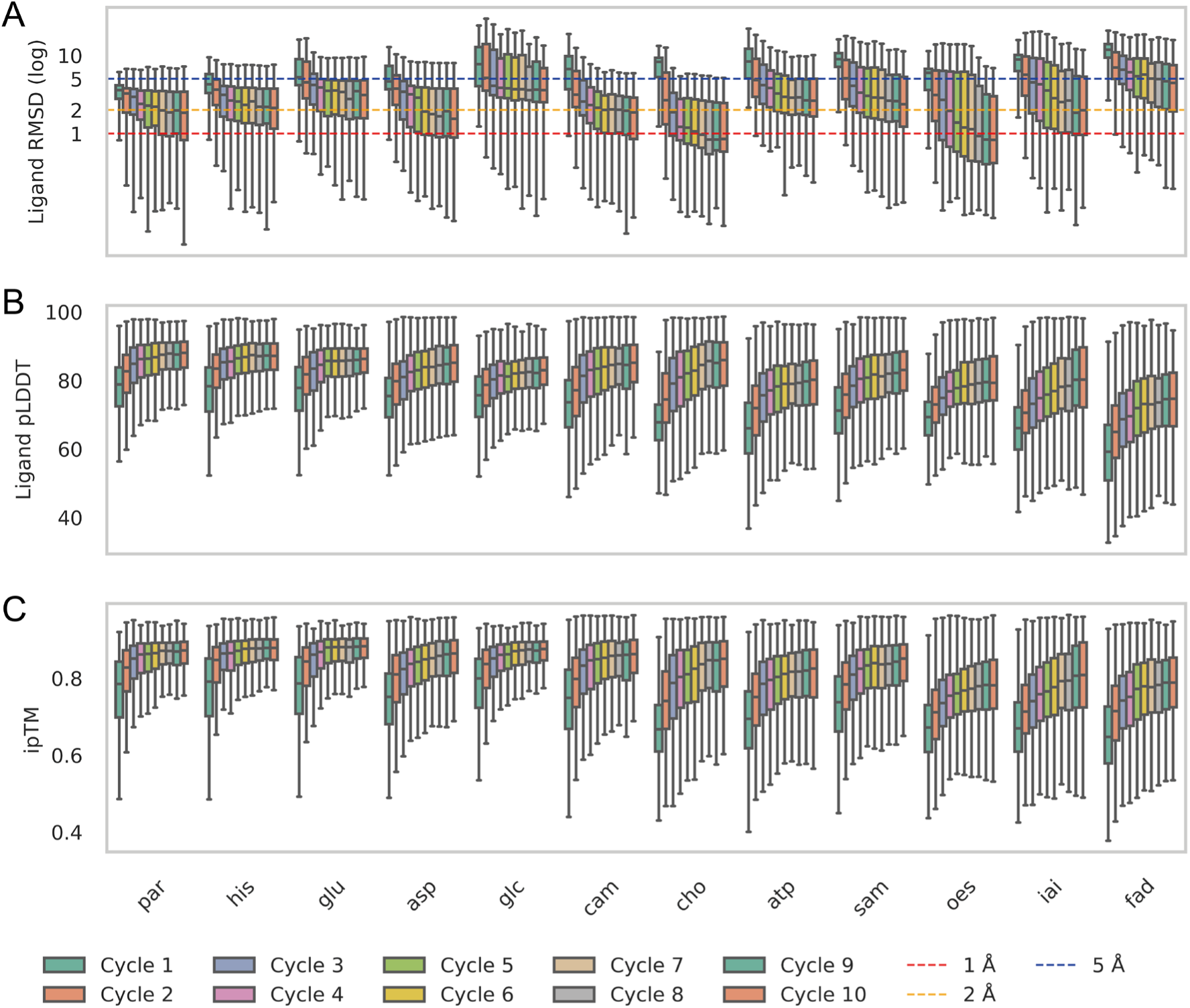
Computation metrics across HalluDesign cycles for black-hole-initialzied small molecule binders. The small molecules are from our in silico dataset shown in **fig. S8**. For each case, the ligand was initially placed randomly at the center of the NTF2 protein scaffold prior to AF3-based HalluDesign optimization. In each cycle, the ligand RMSD, pLDDT, and ipTM distributions of four sequences were evaluated using the AF3 model.

**Fig. S14.**
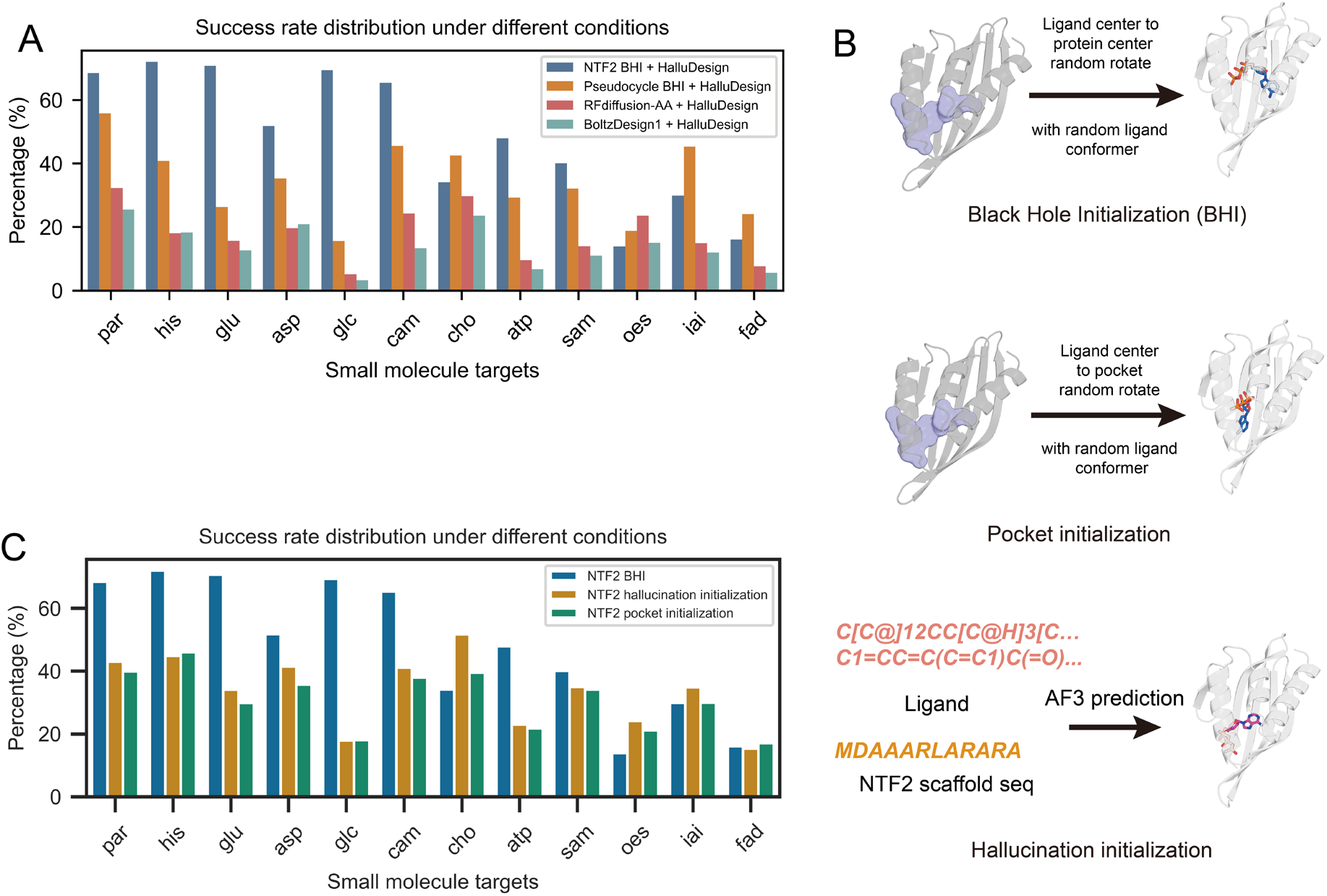
Scaffold-based small molecule binder design. (**A**) Success rates of various small molecule binder design methods. The small molecules are from our in silico dataset shown in **fig. S8**. Initial protein-ligand structures were generated by black-hole-initialization (BHI) using NTF2 or pseudocycle scaffolds, or de novo generated by RFdiffusionAA or BoltzDesign1. HalluDesign were then applied to generated the final structures. Seven optimization cycles were conducted, with truncated diffusion trajectories of 150 steps out of 200 total steps. (**B**) Illustration of the three small molecule initialization approaches. Protein center indicates the center-of-mass of the protein Cα atoms and pocket center is determined by Fpocket (see methods). (**C**) Effect of different initialization strategies (shown in **B**) on final success rates. Final computational metrics were evaluated using the AF3 model. A design was considered successful if it satisfied the following criteria: protein Cα scRMSD ≤ 1.5 Å, protein pLDDT ≥ 85, ligand RMSD ≤ 2 Å, ligand pLDDT ≥ 85, and ipTM ≥ 0.85.

**Fig. S15.**
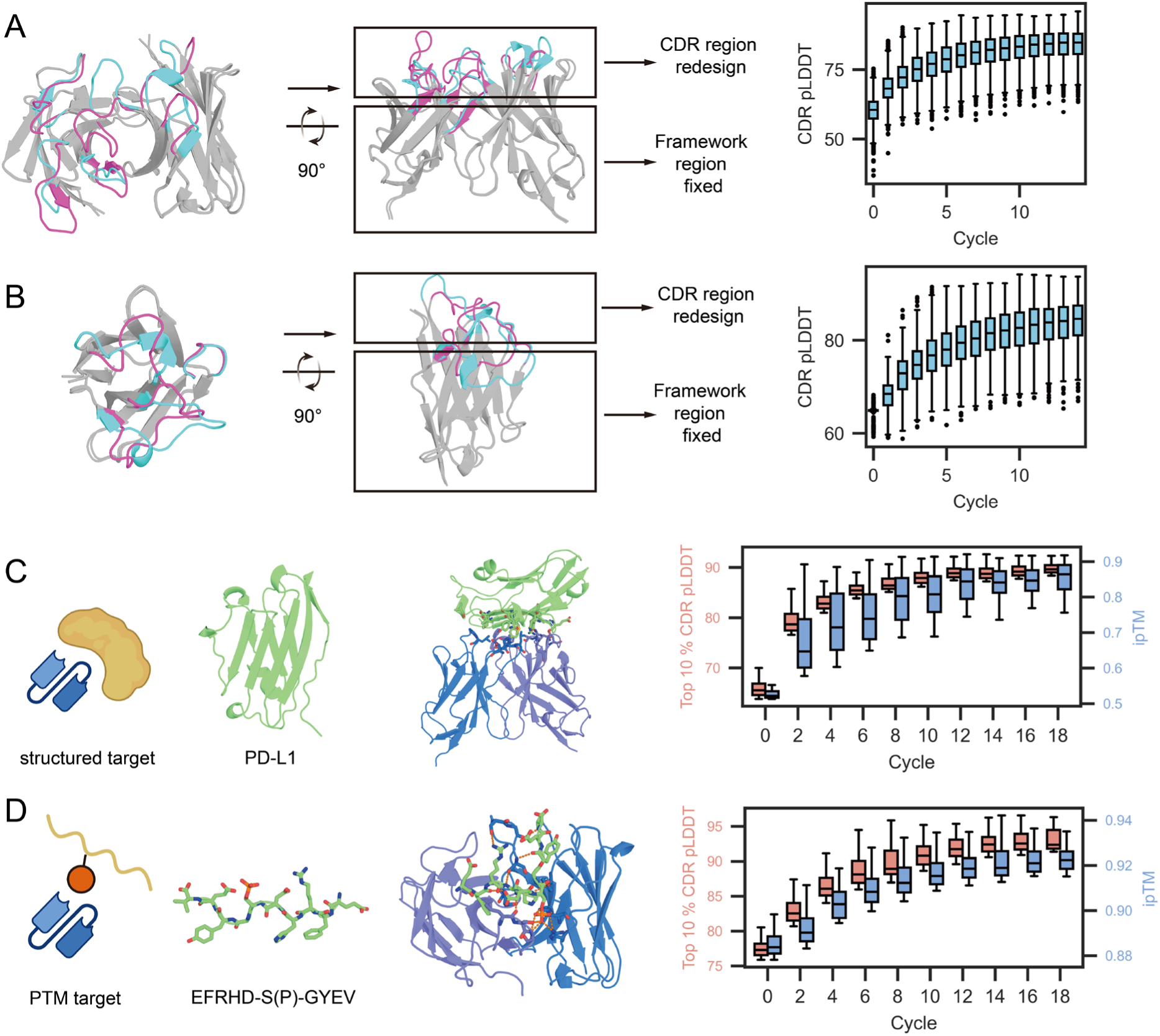
HalluDesign for antibody and nanobody design. (**A, B**) CDR loop redesign in the context of fixed antibody and nanobody frameworks. For antibodies (**A**) and nanobodies (**B**), complementarity-determining regions (CDRs) were iteratively redesigned while maintaining the fixed framework backbones (shown in grey). Superposed structures from the origin CDR region (pink) and the redesigned CDR region (cyan) highlights the changes in CDR loops. Right panels: AF3-predicted pLDDT distributions for redesigned CDR residues across HalluDesign, showing steady improvements in local structural confidence. (**C**) Target specific (PD-L1) antibody design. Left: schematic of the design process. Middle: representative model of designed antibody bound to PD-L1. Right: distribution of pLDDT and ipTM scores for the top 10% highest-scoring designs across 18 HalluDesign optimization cycles. (**D**) Antibody design targeting the phosphorylated Aβ peptide. Left: illustration of the design processes and the phosphorylated Aβ sequence; Middle: representative complex structure. Right: pLDDT and ipTM score distributions for the top 10% designs across 18 HalluDesign cycles.

**Fig. S16.**
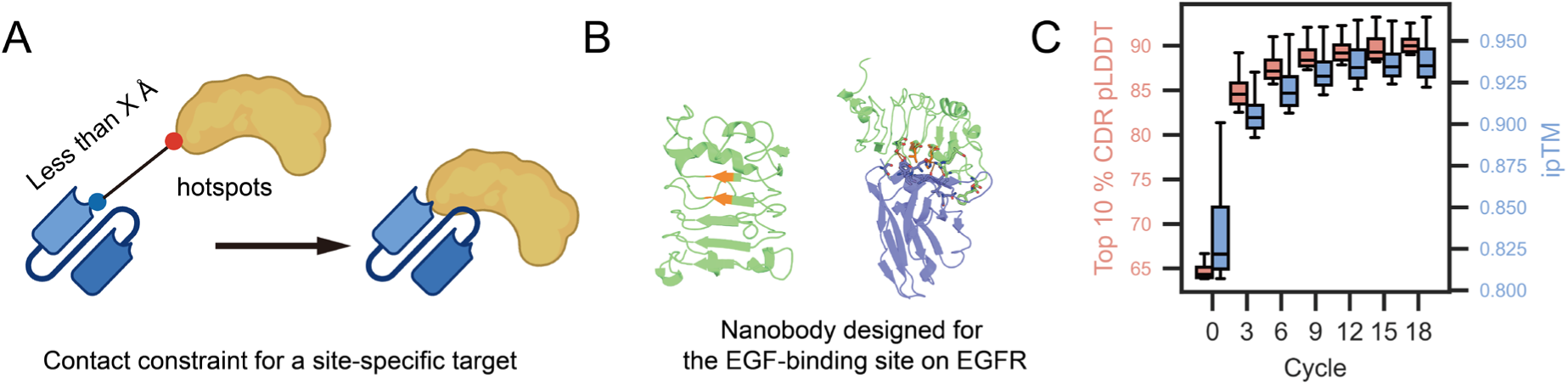
Protenix-based HalluDesign for epitope specific nanobody design. (**A**) HalluDesign utilize contact distance constraint in Protenix as an interaction guidance between CDR and target hotspot residues. (**B**) Representative model of a HalluDesign-generated nanobody with the interaction guidance to target the EGF binding site of EGFR (orange colored). (**C**) Computational metrics across different HalluDesign cycles on epitope specific nanobody design shown in (**B**). The computational metrics were evaluated using the Protenix model.

**Fig. S17.**
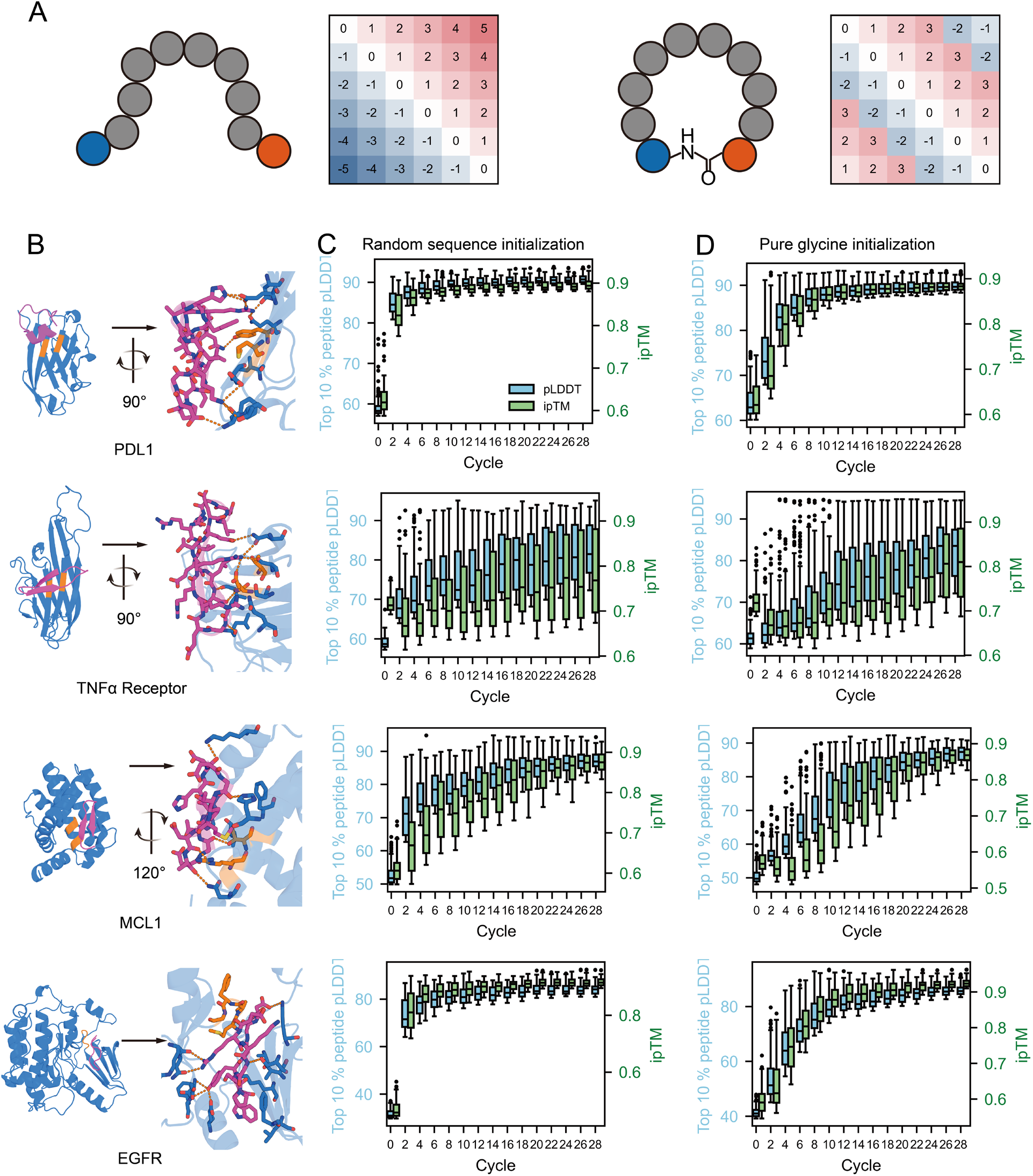
HalluDesign for cyclic peptide binder design. (**A**) Schematic of the positional encoding strategy for cyclic peptide design. Cyclic position encoding is applied to head-to-tail peptides (right), while the default AF3 encoding is linear (left) and generates only linear peptides. **(B)** Representative cyclic peptide binders generated by HalluDesign targeting four different protein targets. (**C, D**) Two sequence initialization approaches were evaluated: random-sequence initialization (**C**) and pure-glycine initialization (**D**). Computation metrics across AF3-based HalluDesign cycles show consistent improvements of model confidence metrics over 30 optimization cycles for both initialization methods (peptide pLDDT in blue, left axis; ipTM in green, right axis). Only the top 10% of highest-confidence sequences generated in each cycle, as evaluated by AF3, are included.

**Fig. S18.**
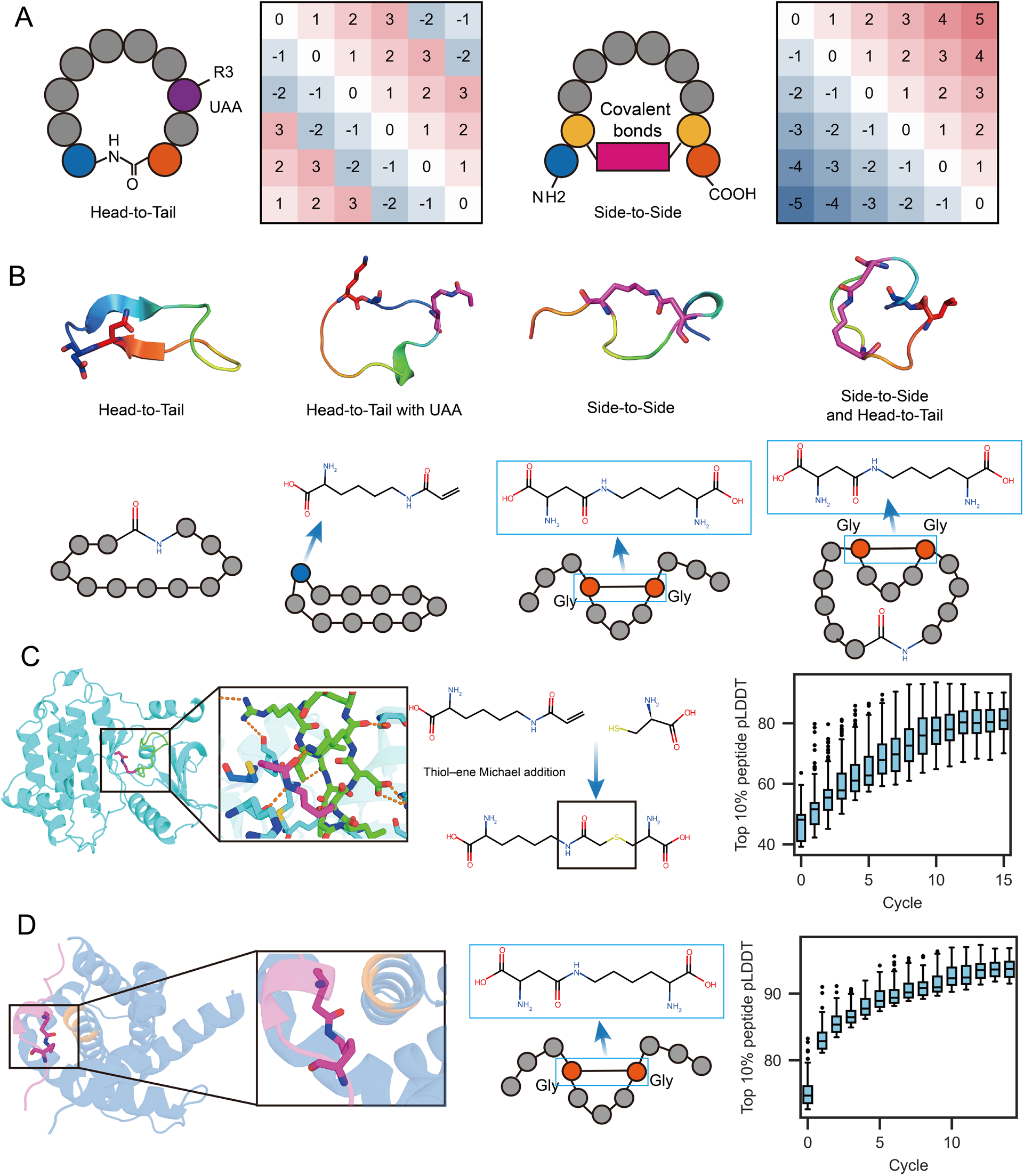
HalluDesign for peptide design featuring diverse cyclic topologies and UAA incorporation. (**A**) Schematic of positional encoding strategies. Cyclic encoding is used for head-to-tail cyclization peptides (left), while the combination of covalent bond definition and linear encoding supports side-to-side architectures (right). (**B**) Representative model structures showing various supported peptide topologies, including standard head-to-tail cyclic peptides, UAA-incorporated cyclic peptides, side-to-side cyclization, and hybrid architectures (detailed implementation is provided in the Supplementary Methods). (**C**) Design of a head-to-tail cyclic peptide intended to covalently bind EGFR Cys797 via UAA ArcK. The inset highlights the covalent attachment. The box plots show the improvement of top 10% pLDDT metric across HalluDesign optimization cycles. (**D**) Design of a side-to-side cyclic peptide targeting MCL1. Structural view (left) and evolution of peptide pLDDT demonstrate successful generation of high-confidence cyclic peptide binder.

**Fig. S19.**
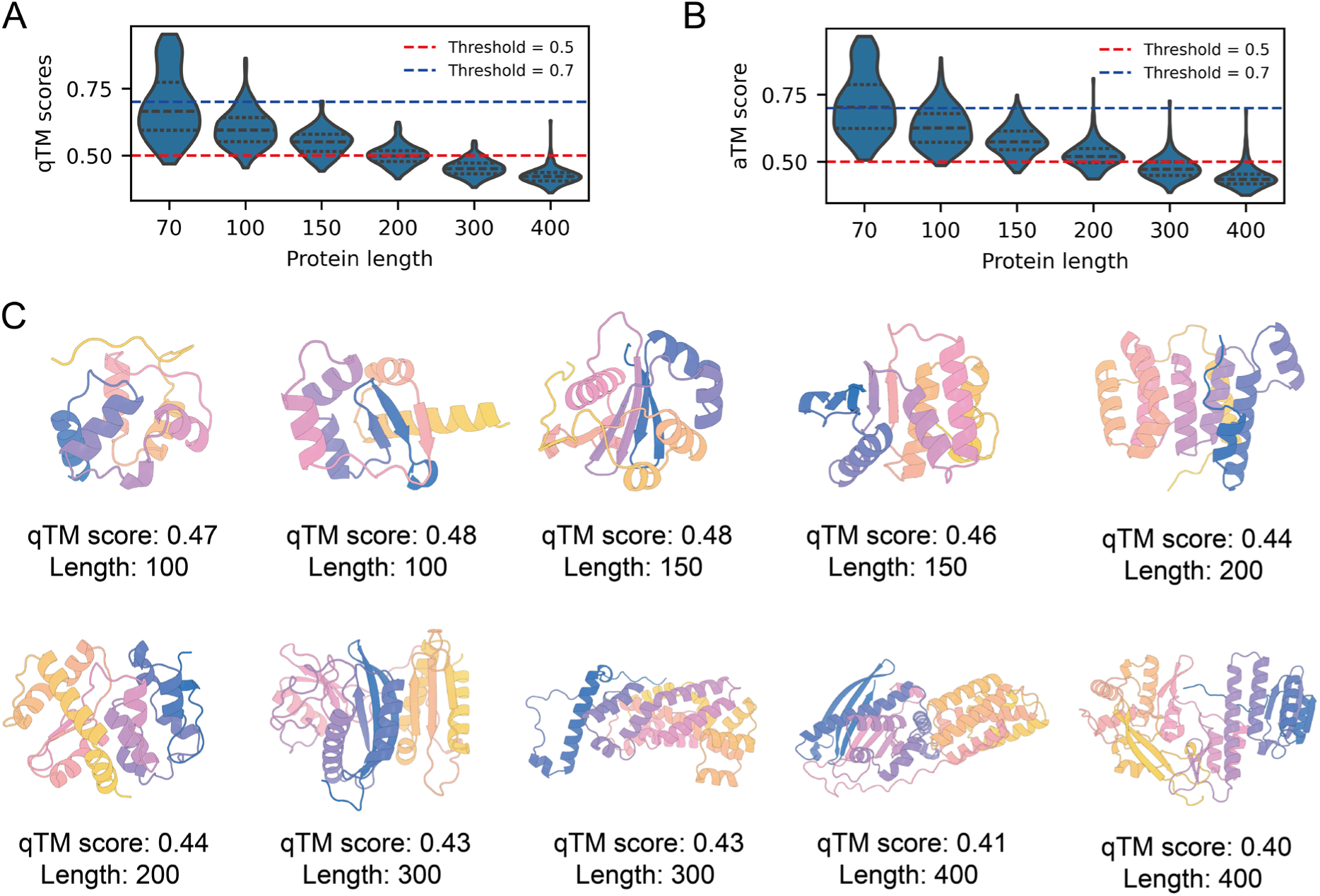
HalluDesign for unconditional protein monomer generation. Distributions of best-hit FoodSeek Query TM scores (**A**) and Alignment TM scores (**B**) for HalluDesign-generated protein monomers of varying lengths (70–400 residues). (**C**) Representative examples and their qTM scores of HalluDesign-generated structures across different sequence lengths.

**Fig. S20.**
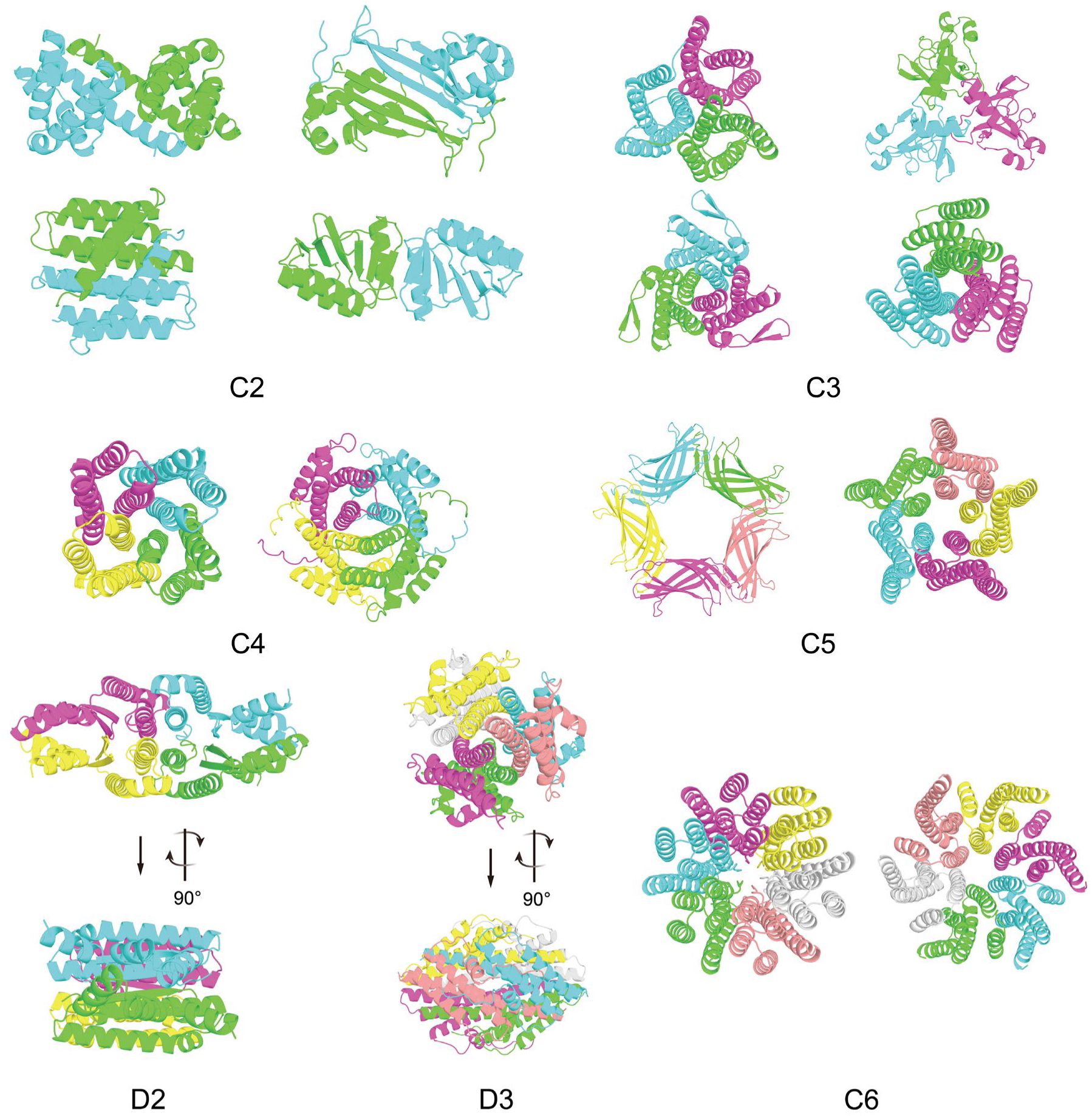
HalluDesign for symmetric protein assemblies design. Representative examples of de novo generated complexes spanning multiple symmetry groups, including cyclic (C2–C6) and dihedral (D2, D3) symmetries. The generated scaffolds exhibit a broad repertoire of topologies, including all-helical bundles, mixed α/β and all β structures.

**Fig. S21.**
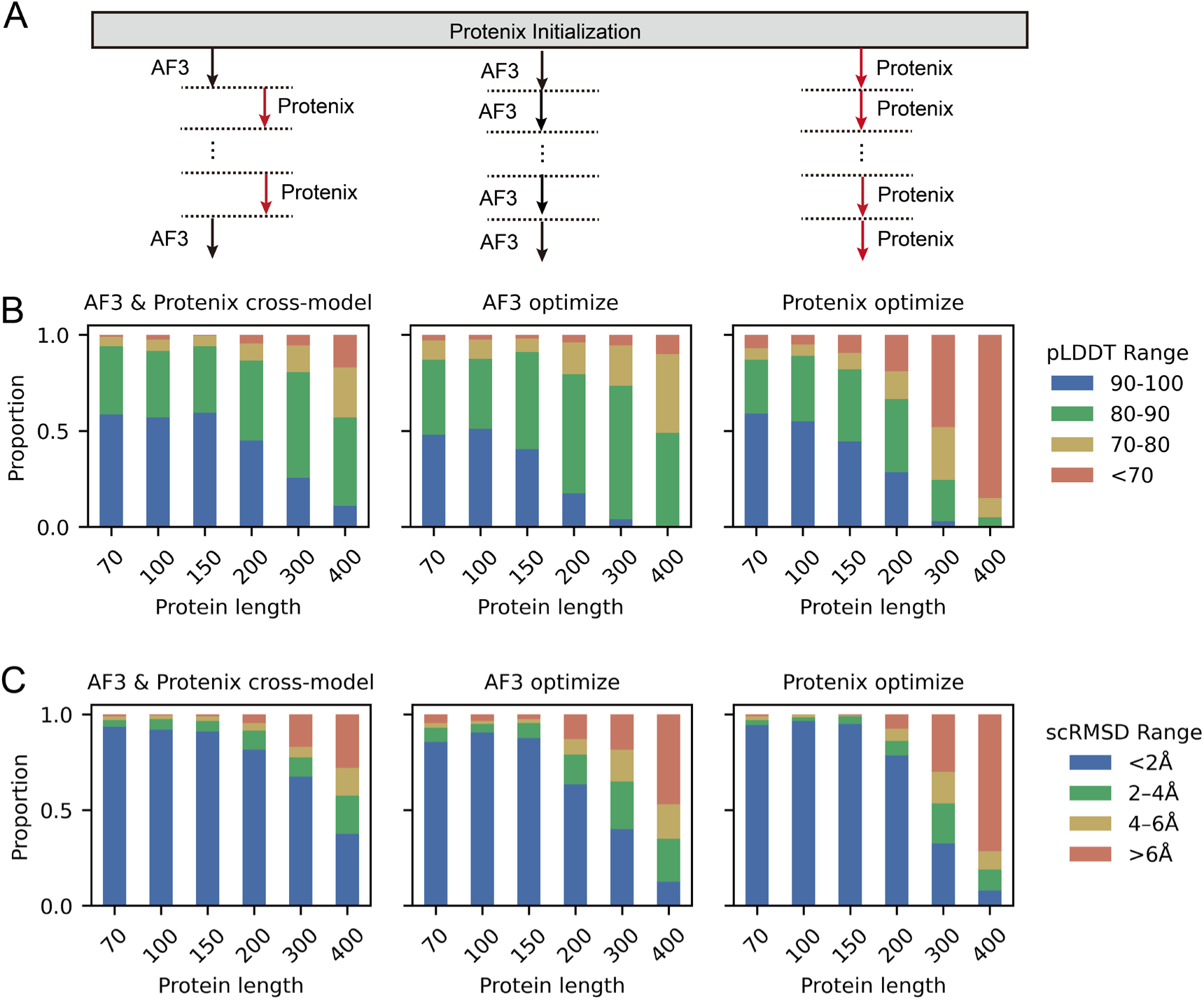
The AF3–Protenix cross-cycle optimization improves protein structure generation quality across various sequence lengths. Quality metrics for proteins ranging from 70 to 400 residues are compared among three optimization strategies: **(A)** Cross-cycle (left), AF3-based (middle), and Protenix-based (right). All designs are initialized by Protenix-based HalluDesign with random sequence. (**B**) Distribution of pLDDT confidence scores, categorized into four ranges. The cross-cycle strategy maintains a higher proportion of high-confidence designs (blue, >90). (**C**) Distribution of Cα RMSD between designed structures and their AF3-predicted models. The cross-cycle approach consistently yields lower RMSD values (<2 Å) compared to single-model optimization.

**Fig. S22.**
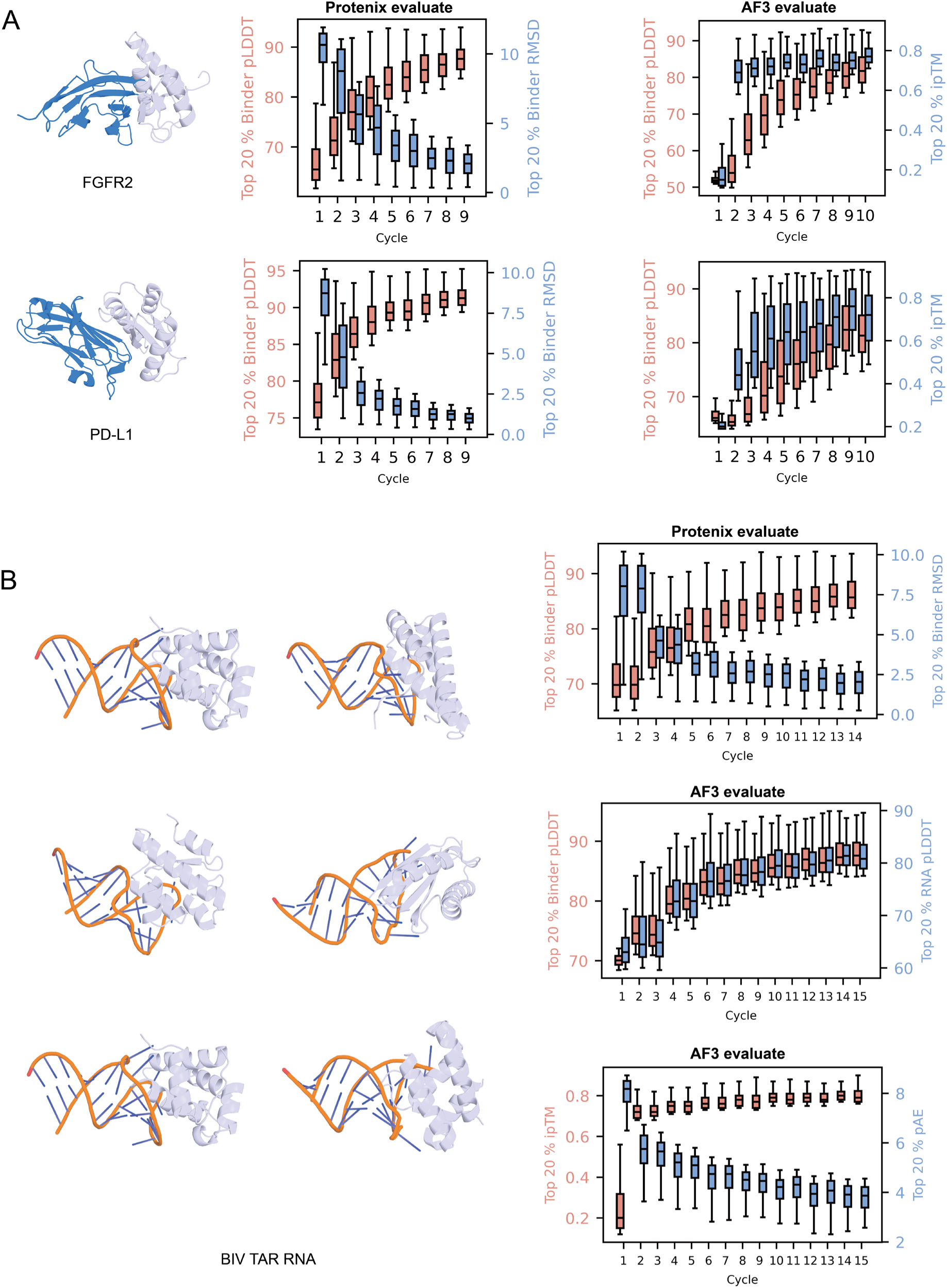
HalluDesign for from scratch binder design targeting protein and RNA. (**A**) Design of protein binders targeting FGFR2 (top row) and PD-L1 (bottom row). Left: Representative structures of the designed binders (light blue) in complex with their respective targets (dark blue). Middle: Protenix-based evaluation tracking the distribution of the top 20% values for binder pLDDT (red) and binder RMSD (blue) independently at each cycle. Right: Validation using AlphaFold3 (AF3), showing the distributions of the top 20% binder pLDDT scores (red) and top 20% ipTM scores (blue). (**B**) Design of RNA binders targeting the BIV TAR RNA. Left: Representative models of designed protein binders (light blue) bound to the structured RNA target (orange backbone with blue bases). Right: Steady improvement of confidence scores across HalluDesign optimization cycles. The box plots display the top 20% distributions for each metric: Protenix-predicted binder pLDDT and RMSD (top), AF3-predicted binder and RNA pLDDT scores (middle), and AF3-predicted complex quality metrics including ipTM and pAE (bottom).

**Fig. S23.**
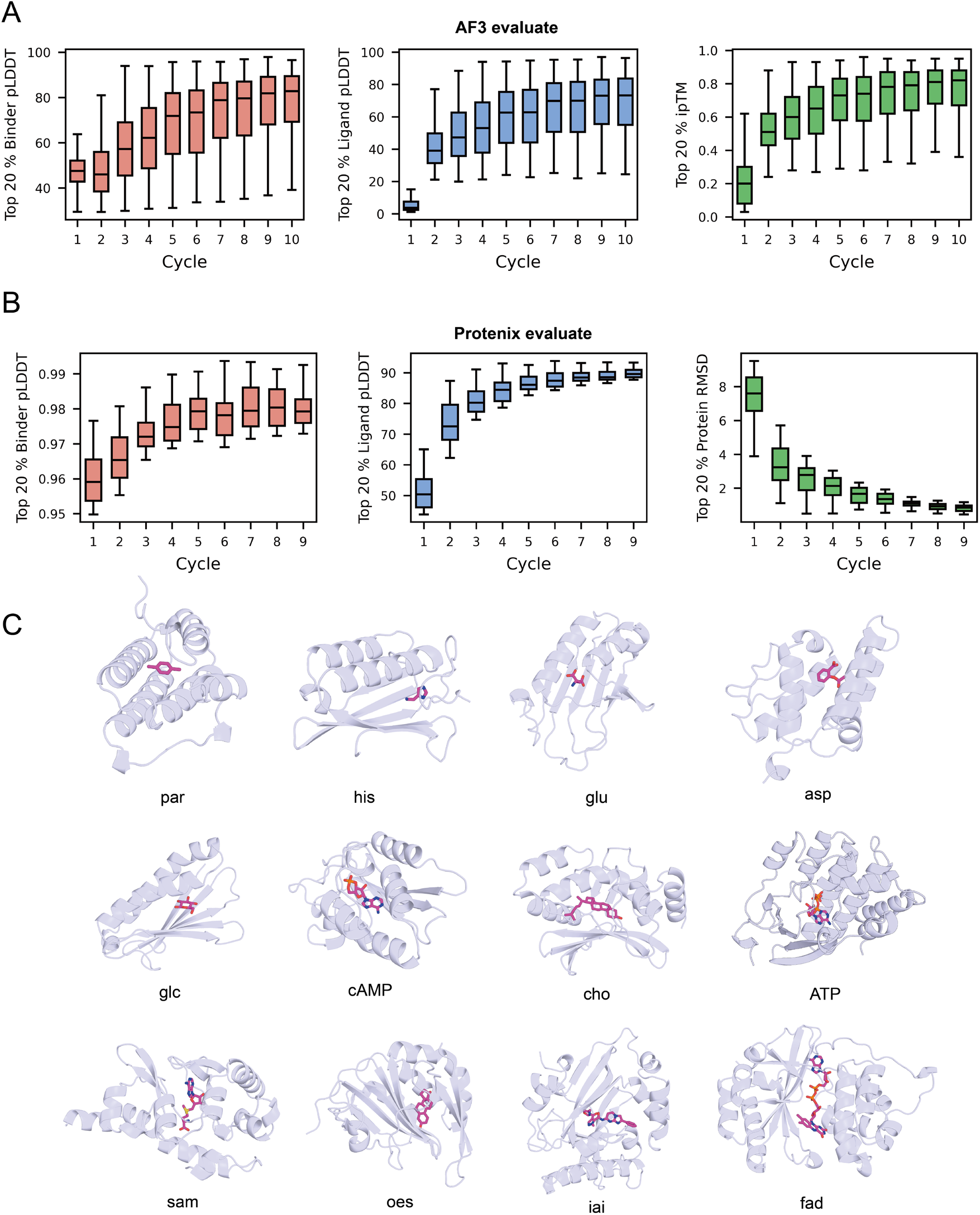
From scratch small molecule binders using HalluDesign. Averaged computational metrics evaluated by AF3 (**A**) and Protenix (**B**) over the small molecule targets in **fig. S8** across the HalluDesign optimization cycles. The plots show the distribution of the top 20% best designs of each metric: binder pLDDT (red), ligand pLDDT (blue), and ipTM (for AF3) or protein scRMSD (for Protenix) (green) at each cycle. (**C**) Representative structural models of designed proteins (light blue) in complex with various small molecules in **fig. S8**.

**Fig. S24.**
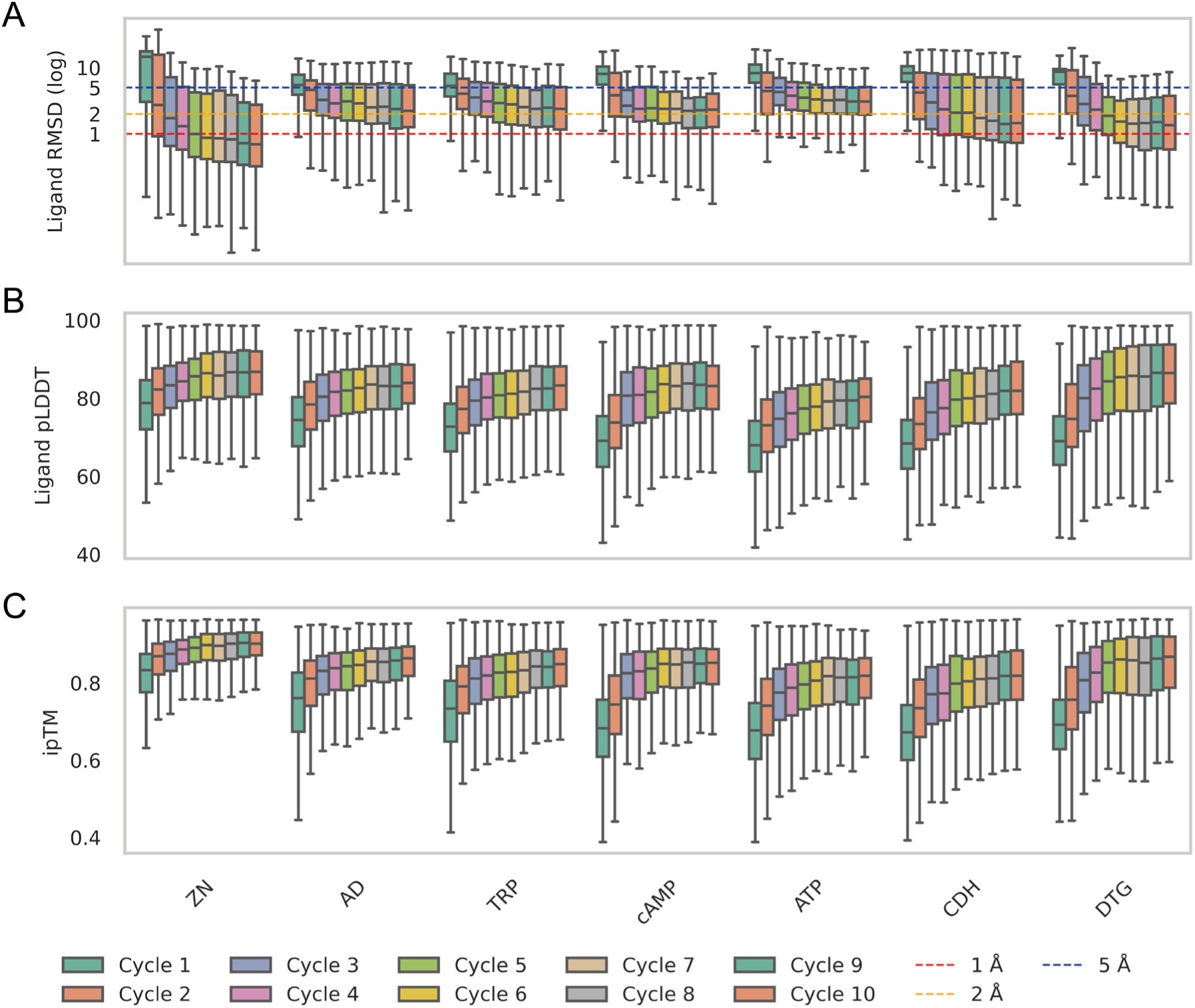
HalluDesign optimization trajectories for the experimentally validated small molecule targets. The initial complex structures were generated using the black-hole initialization strategy, followed by AF3-based HalluDesign optimization to produce the final complex structures. Computational metrics including ligand RMSD, pLDDT, and ipTM were evaluated with AF3 across the HalluDesign optimization cycles.

**Fig. S25.**
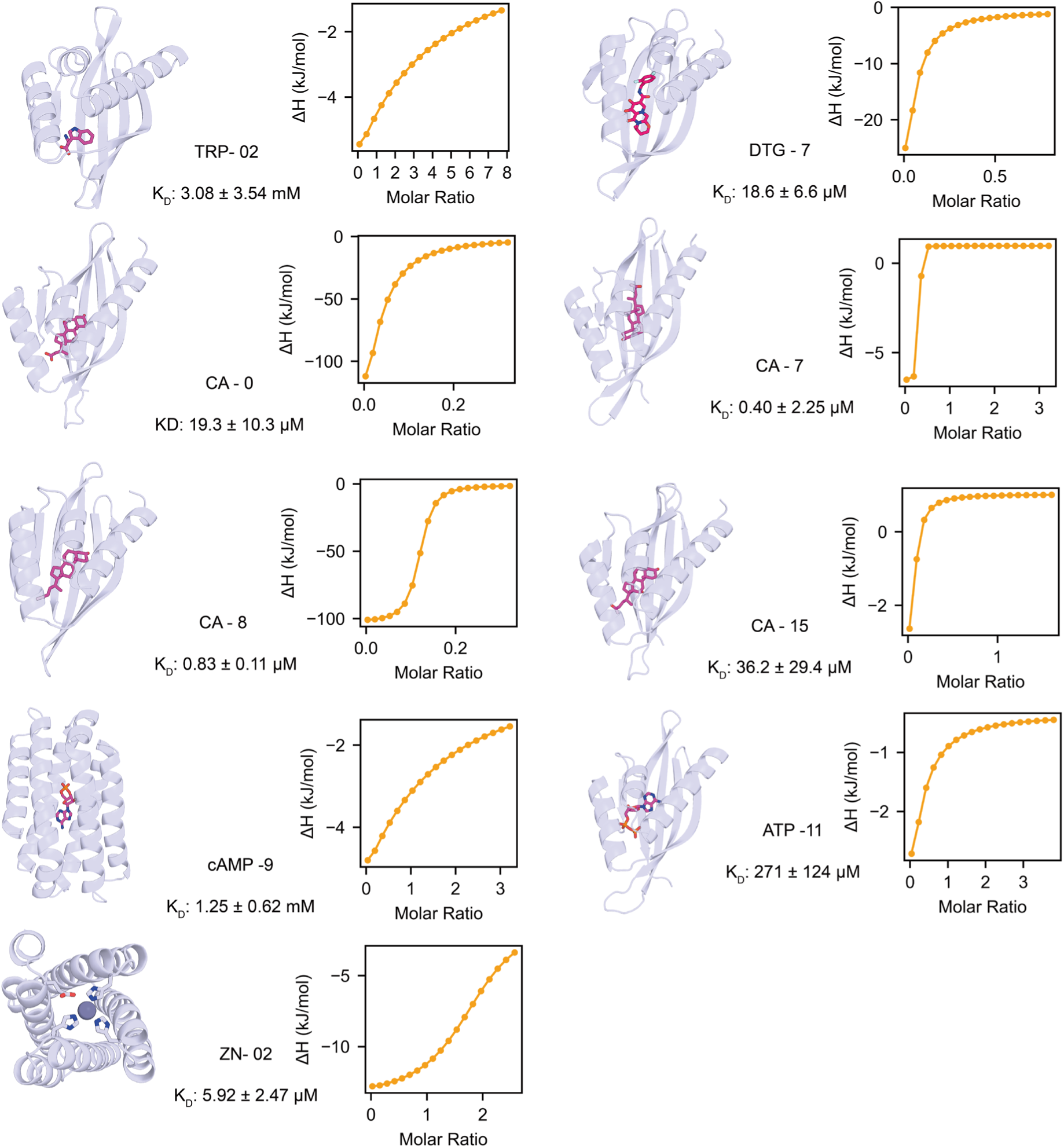
ITC results for HalluDesign-generated ligand binders not shown in Fig. 4.

**Fig. S26.**
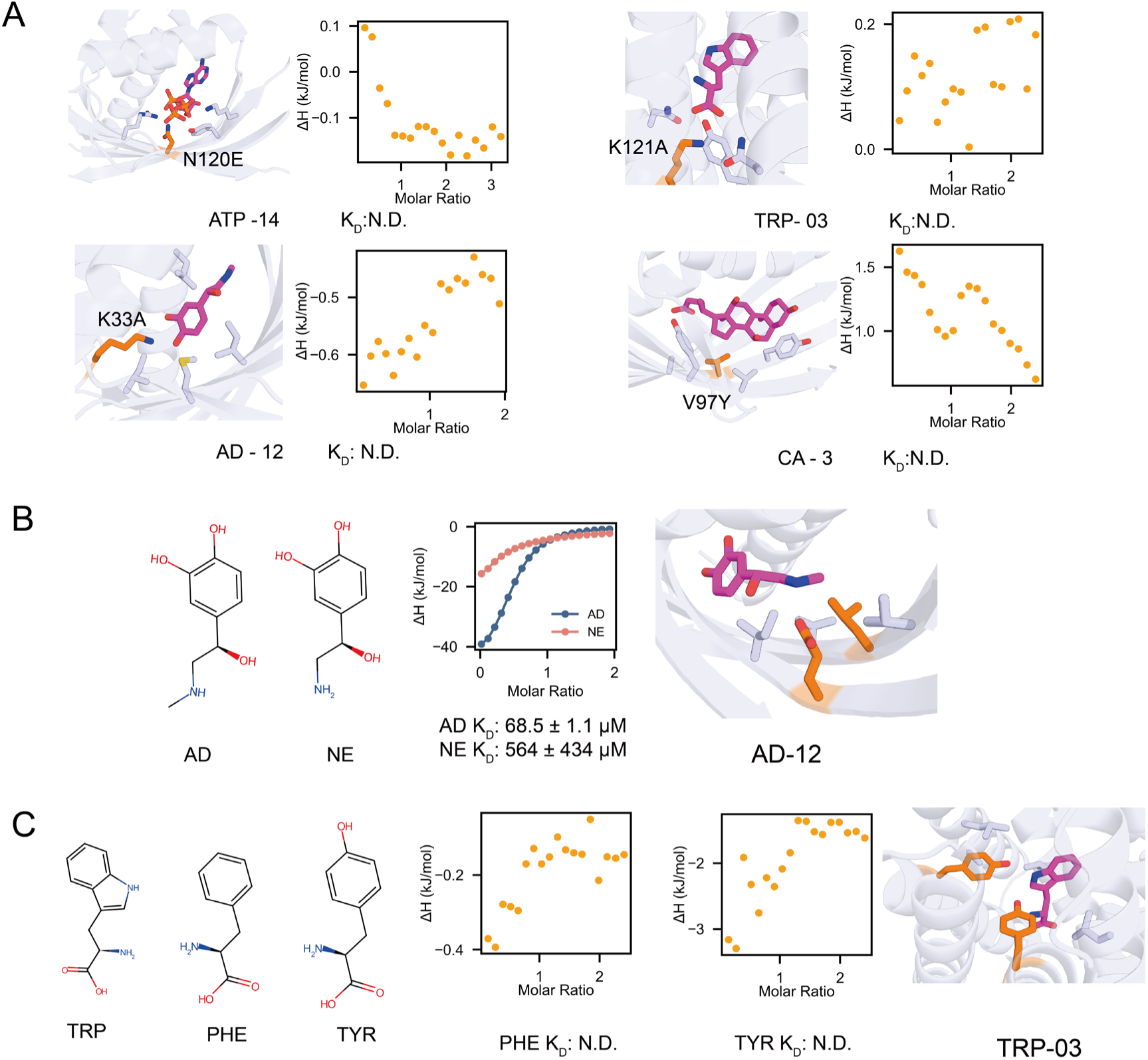
Experimental characterization of binding specificity and mutational analysis of ligand-binding proteins. (**A**) Zoomed in views of ligand binding pockets highlighting mutated residues (orange) and the corresponding ITC results for these mutants. (**B**) Ligand binding specificity of the AD-12 binder. The design differentiates adrenaline (AD) from noradrenaline (NE), exhibiting strong selectivity with approximately eightfold higher affinity for AD. (**C**) Ligand specificity of the TRP-03 binder. The design binds specifically to tryptophan, with no detectable binding to phenylalanine (PHE) and tyrosine (TYR).

**Fig. S27.**
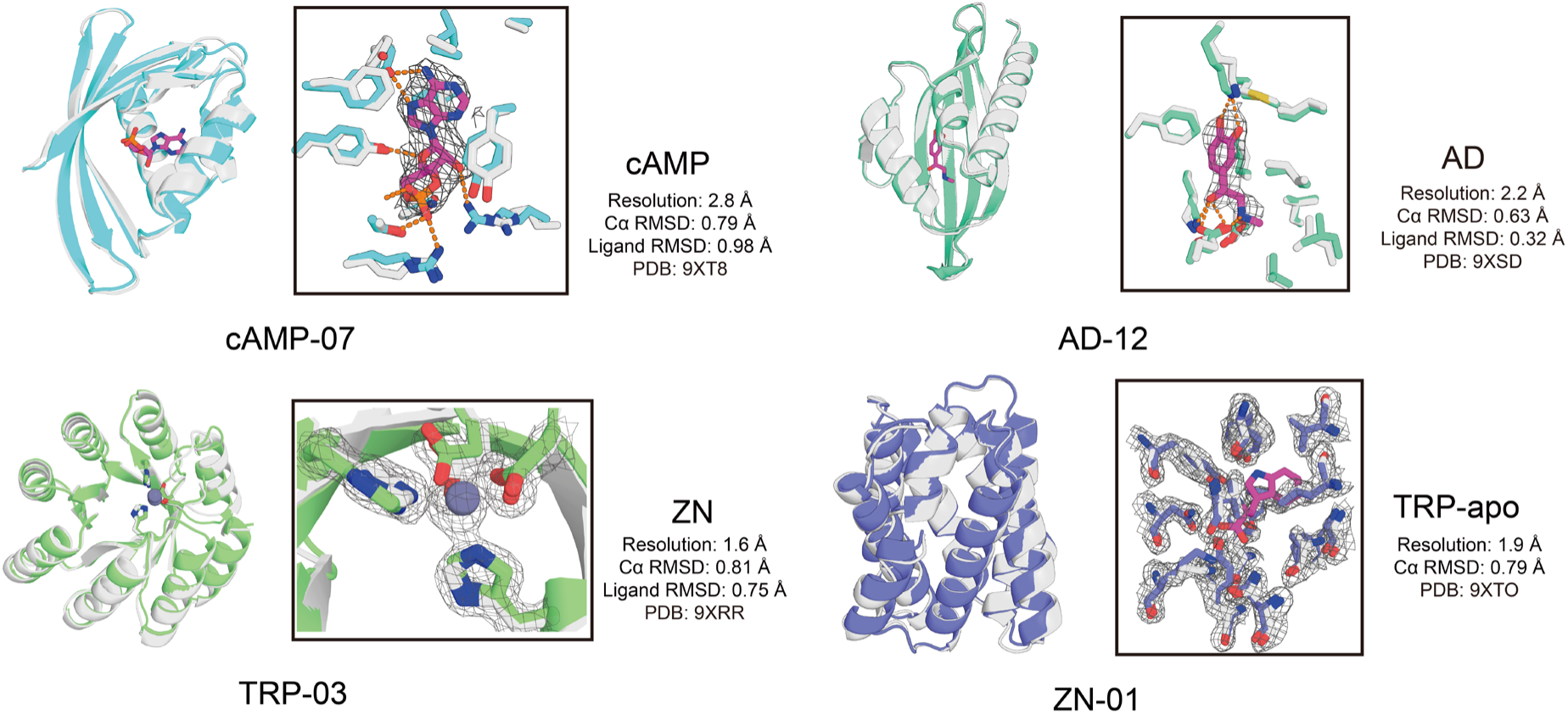
Crystal structure validation of HalluDesign-generated small molecule binders. The design model is shown in white, overlaid with the experimentally determined crystal structure and electron density map. The composite omit electron density for the ligand and neighboring residues is contoured at 1.8σ.

**Fig. S28.**
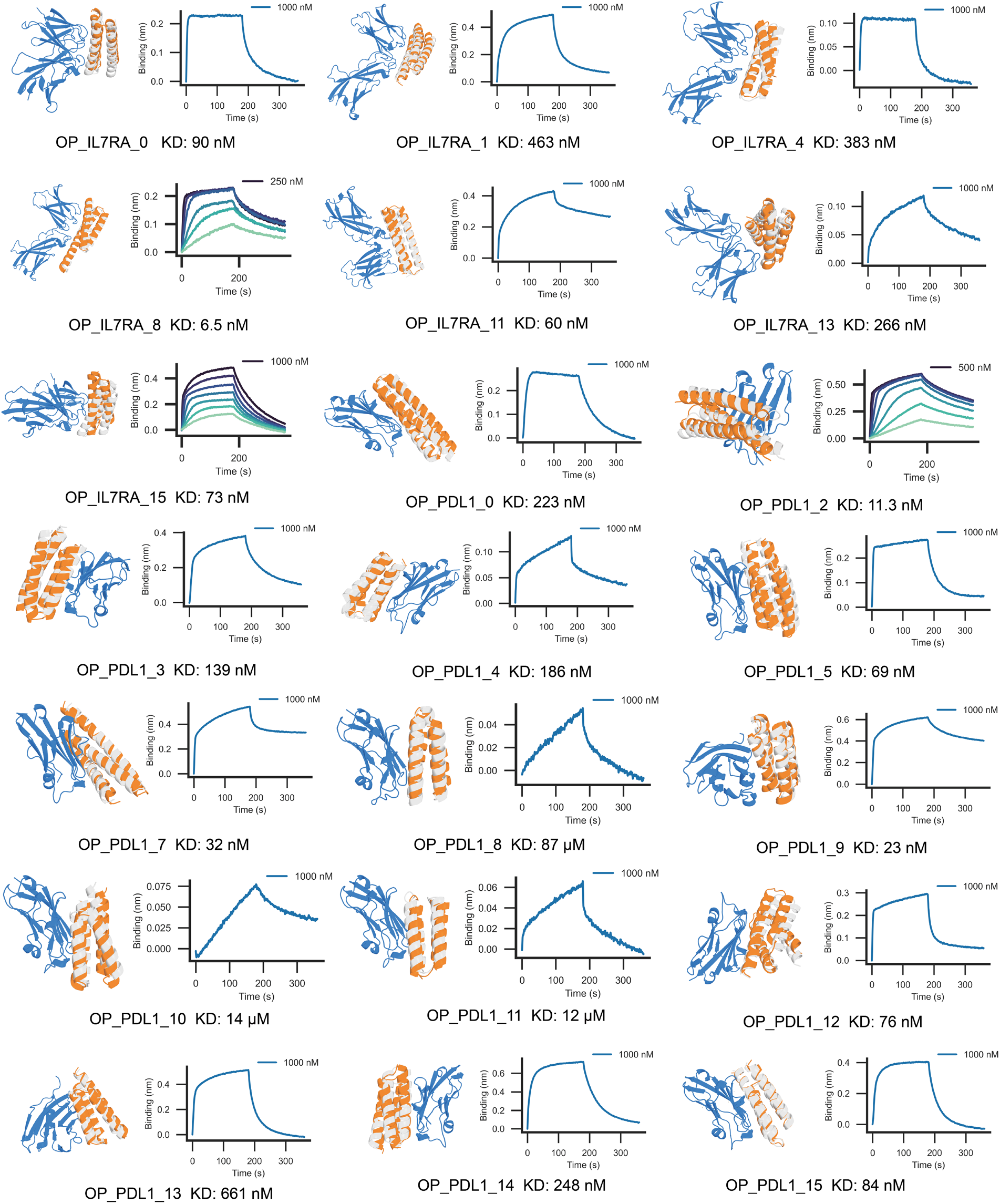
BLI binding analysis of rescued RFdiffusion binders. For each target, one or more high-affinity binders identified during initial screening were selected for BLI titration analysis for more accurate binding affinity determination. For the shown design models, orange represents the HalluDesign-optimized structure, white the original RFdiffusion structure, and blue the target protein. For BLI titration experiments, only the sensorgram at the highest ligand concentration is labeled; the remaining concentrations follow a two-fold serial dilution.

**Fig. S29.**
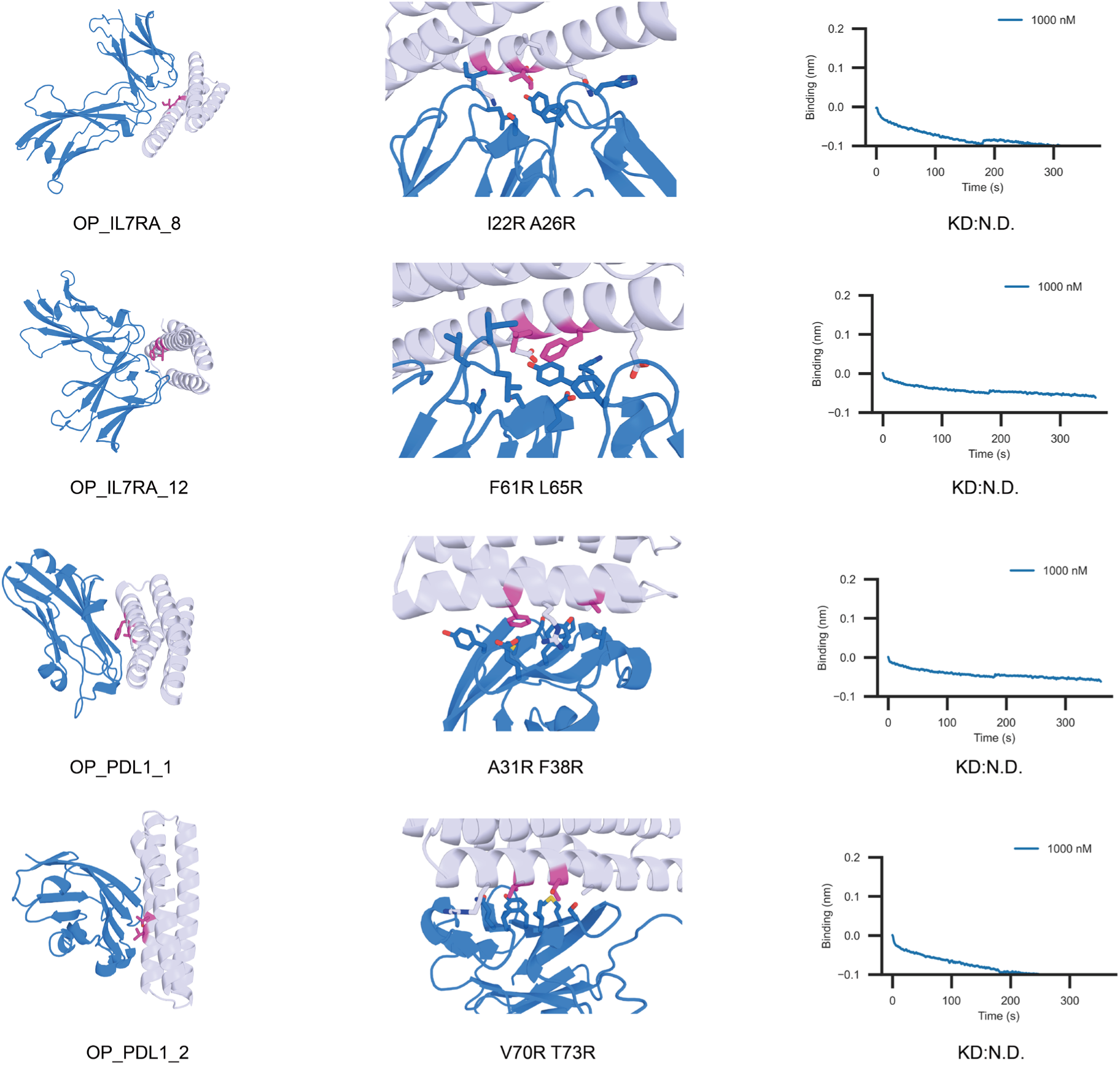
Interface mutational analysis of rescued RFdiffusion binders. N.D. indicates no detectable binding. Each row displays the overall structure (left), a zoomed-in view of the interface with mutated residues highlighted in pink (center), and the corresponding BLI binding curves (right).

**Fig. S30.**
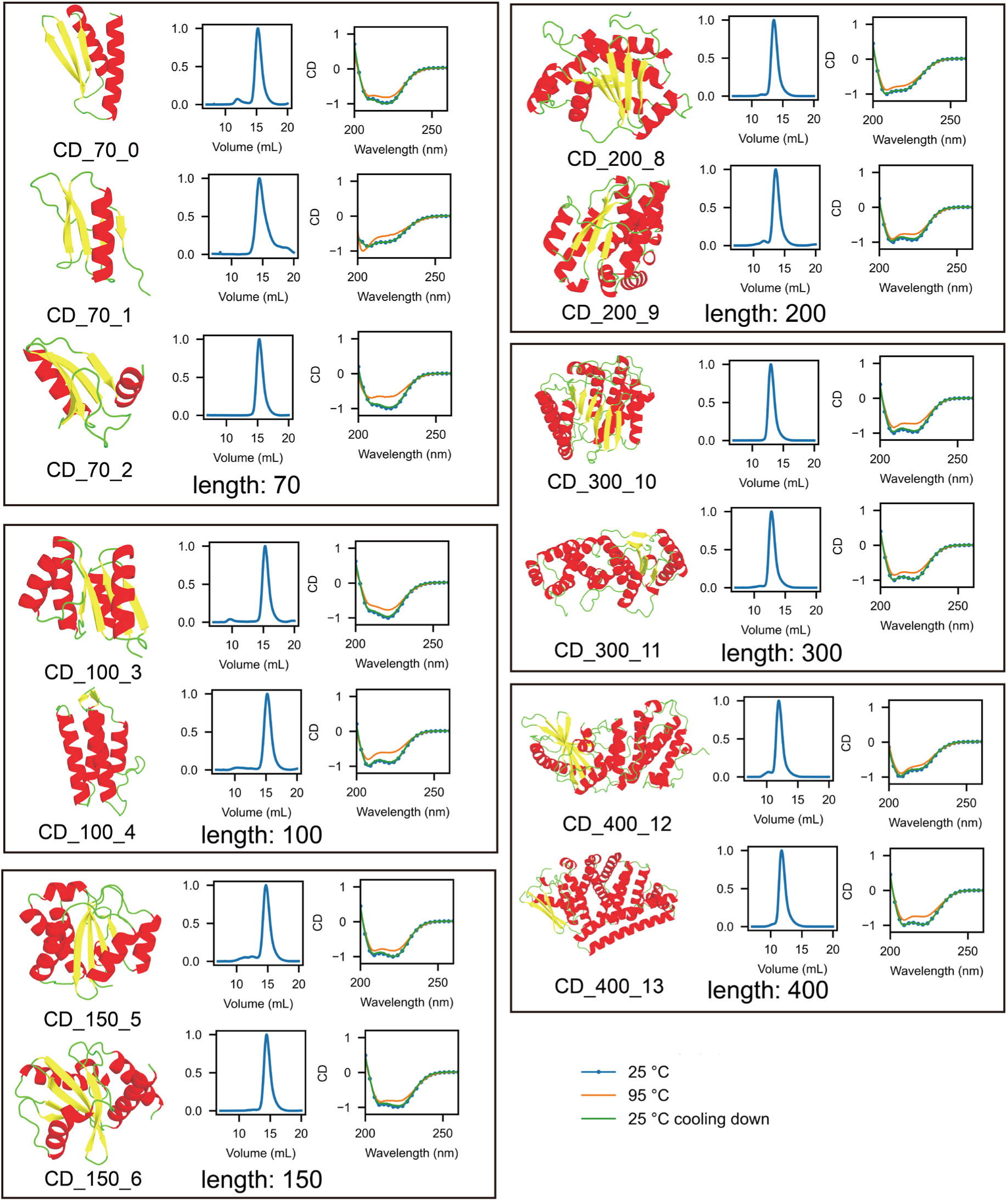
Biophysical characterization of HalluDesign-generated protein monomers across varying sequence lengths. Designs are categorized by length (70 to 400 residues). Panels display the design model (left), SEC elution profile (middle), and CD spectra (right) for each design. SEC data confirm monodispersity, while CD spectras at 25°C, 95°C, and upon refolding (cooling) verify that the proteins fold into stable structures.

**Fig. S31.**
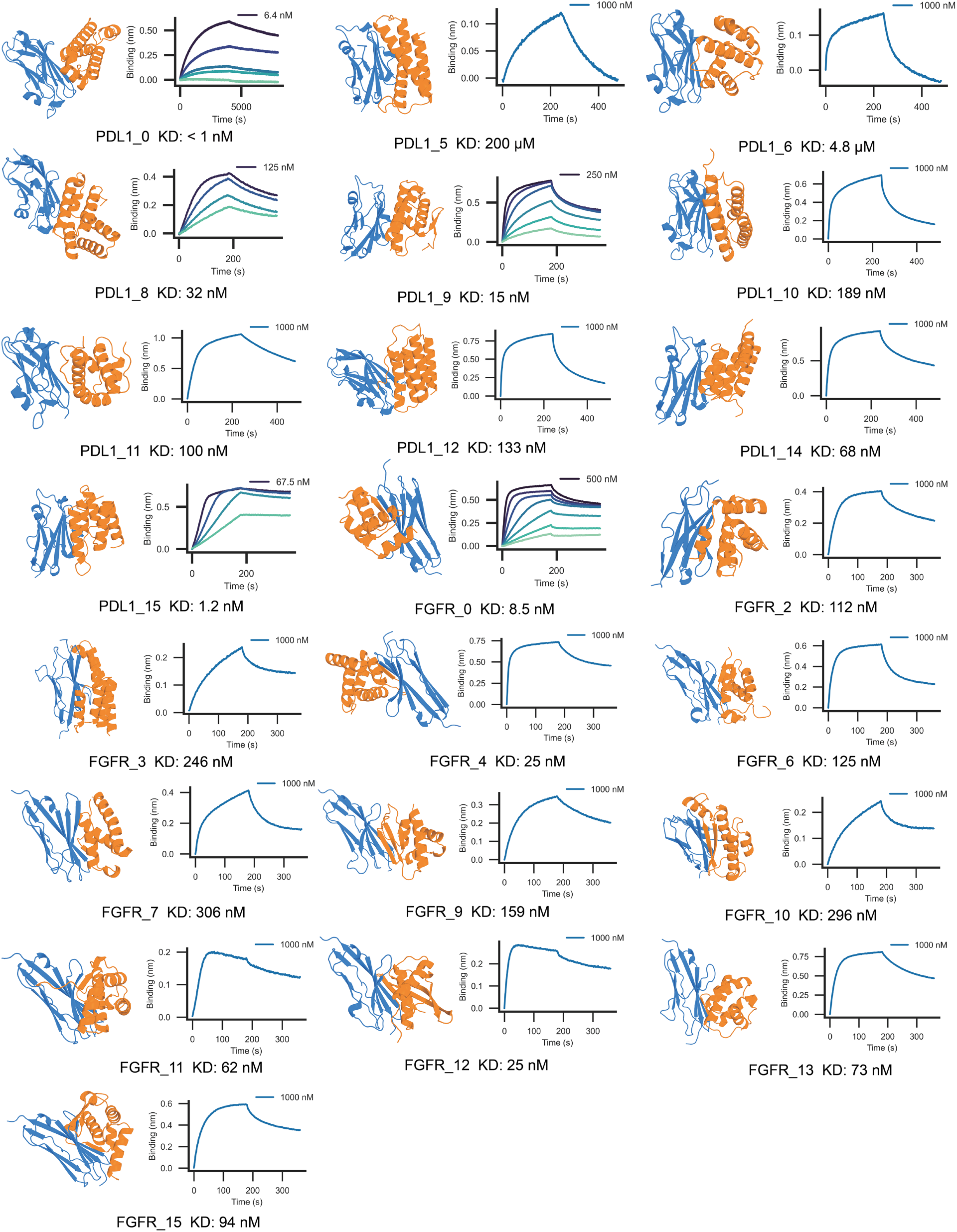
BLI binding analysis of the from scratch protein binders generated by HalluDesign. For each target, one or more high-affinity binders identified during initial screening were selected for BLI titration analysis for more accurate binding affinity determination. For the shown design model, orange represents the bind and blue the target protein. For BLI titration experiments, only the sensorgram at the highest ligand concentration is labeled; the remaining concentrations follow a two-fold serial dilution.

**Fig. S32.**
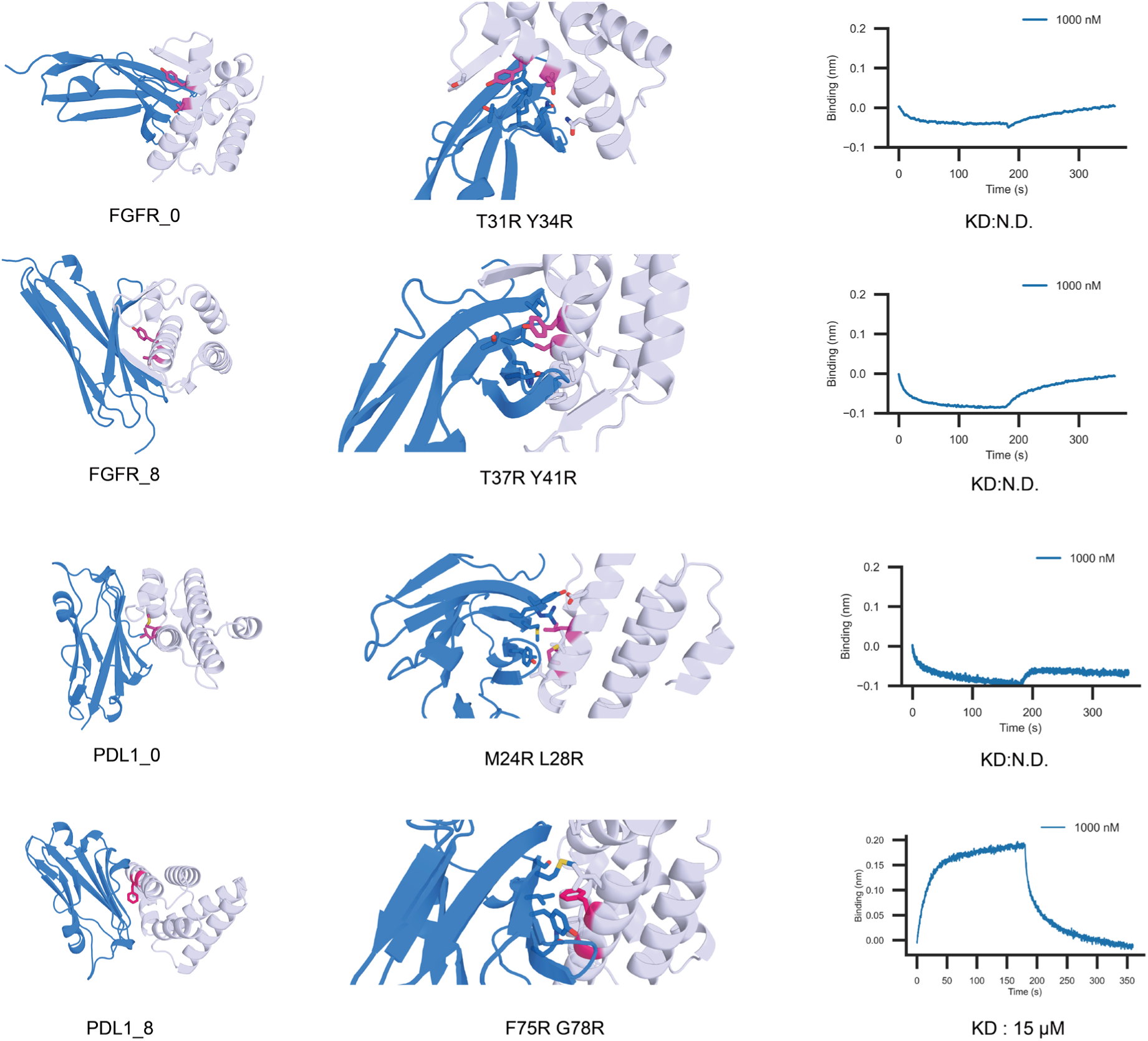
Interface mutational analysis of from scratch designed protein binders. N.D. indicates no detectable binding. Each row includes the overall structure (left), a zoomed-in view of the interface with mutated residues highlighted in pink (center), and the BLI binding curve of the corresponding mutant (right).

## References and Notes

1. Z. Du et al., The trRosetta server for fast and accurate protein structure prediction. Nature protocols 16, 5634–5651 (2021).

2. R. Evans et al., Protein complex prediction with AlphaFold-Multimer. biorxiv, 2021.2010. 2004.463034 (2021).

3. J. Jumper et al., Highly accurate protein structure prediction with AlphaFold. nature 596, 583–589 (2021).

4. Z. Lin et al., Evolutionary-scale prediction of atomic-level protein structure with a language model. Science 379, 1123–1130 (2023).

5. I. Anishchenko et al., De novo protein design by deep network hallucination. Nature 600, 547–552 (2021).

6. B. I. Wicky et al., Hallucinating symmetric protein assemblies. Science 378, 56–61 (2022).

7. C. Norn et al., Protein sequence design by conformational landscape optimization. Proceedings of the National Academy of Sciences 118, e2017228118 (2021).

8. C. Frank, D. Schiwietz, L. Fuß, S. Ovchinnikov, H. Dietz, Alphafold2 refinement improves designability of large de novo proteins. bioRxiv, 2024.2011. 2021.624687 (2024).

9. M. Pacesa et al., BindCraft: one-shot design of functional protein binders. bioRxiv, 2024.2009. 2030.615802 (2024).

10. A. H.-W. Yeh et al., De novo design of luciferases using deep learning. Nature 614, 774–780 (2023).

11. Y. Liu et al., De novo protein design with a denoising diffusion network independent of pretrained structure prediction models. Nature Methods 21, 2107–2116 (2024).

12. C. Wang et al., Proteus: exploring protein structure generation for enhanced designability and efficiency. bioRxiv, 2024.2002. 2010.579791 (2024).

13. N. R. Bennett et al., Atomically accurate de novo design of antibodies with RFdiffusion. Nature, 1–11 (2025).

14. C. Liu et al., Diffusing protein binders to intrinsically disordered proteins. Nature, 1–9 (2025).

15. J. Wang et al., Scaffolding protein functional sites using deep learning. Science 377, 387–394 (2022).

16. J. Abramson et al., Accurate structure prediction of biomolecular interactions with AlphaFold 3. Nature 630, 493–500 (2024).

17. D. Desai et al., Review of AlphaFold 3: transformative advances in drug design and therapeutics. Cureus 16, (2024).

18. C. D. team et al., Chai-1: Decoding the molecular interactions of life. BioRxiv, 2024.2010. 2010.615955 (2024).

19. B. A. A. S. Team et al., Protenix-advancing structure prediction through a comprehensive AlphaFold3 reproduction. BioRxiv, 2025.2001. 2008.631967 (2025).

20. J. Wohlwend et al., Boltz-1 democratizing biomolecular interaction modeling. BioRxiv, 2024.2011. 2019.624167 (2025).

21. Y. Cho, M. Pacesa, Z. Zhang, B. Correia, S. Ovchinnikov, BoltzDesign1: Inverting All-Atom Structure Prediction Model for Generalized Biomolecular Binder Design. bioRxiv, 2025.2004. 2006.647261 (2025).

22. M. Ren et al., PXDesign: Fast, Modular, and Accurate De Novo Design of Protein Binders. bioRxiv, 2025.2008. 2015.670450 (2025).

23. C. D. Team et al., Zero-shot antibody design in a 24-well plate. bioRxiv, 2025.2007. 2005.663018 (2025).

24. O. Zhang, et al., ODesign: A World Model for Biomolecular Interaction Design. arXiv preprint arXiv:2510.22304, (2025).

25. A. Leaver-Fay et al., in Methods in enzymology. (Elsevier, 2013), vol. 523, pp. 109–143.

26. R. F. Alford et al., The Rosetta all-atom energy function for macromolecular modeling and design. Journal of chemical theory and computation 13, 3031–3048 (2017).

27. B. Kuhlman, P. Bradley, Advances in protein structure prediction and design. Nature reviews molecular cell biology 20, 681–697 (2019).

28. I. V. Korendovych, W. F. DeGrado, De novo protein design, a retrospective. Quarterly reviews of biophysics 53, e3 (2020).

29. J. Dauparas et al., Robust deep learning–based protein sequence design using ProteinMPNN. Science 378, 49–56 (2022).

30. J. Dauparas et al., Atomic context-conditioned protein sequence design using LigandMPNN. Nature Methods, 1–7 (2025).

31. R. Hadsell, S. Chopra, Y. LeCun, in 2006 IEEE computer society conference on computer vision and pattern recognition (CVPR’06). (IEEE, 2006), vol. 2, pp. 1735–1742.

32. M. Varadi et al., AlphaFold Protein Structure Database: massively expanding the structural coverage of protein-sequence space with high-accuracy models. Nucleic acids research 50, D439–D444 (2022).

33. J. B. Ingraham et al., Illuminating protein space with a programmable generative model. Nature 623, 1070–1078 (2023).

34. C. Frank et al., Scalable protein design using optimization in a relaxed sequence space. Science 386, 439–445 (2024).

35. J. L. Watson et al., De novo design of protein structure and function with RFdiffusion. Nature 620, 1089–1100 (2023).

36. R. Krishna et al., Generalized biomolecular modeling and design with RoseTTAFold All-Atom. Science 384, eadl2528 (2024).

37. B. Basanta et al., An enumerative algorithm for de novo design of proteins with diverse pocket structures. Proceedings of the National Academy of Sciences 117, 22135–22145 (2020).

38. L. An et al., Binding and sensing diverse small molecules using shape-complementary pseudocycles. Science 385, 276–282 (2024).

39. B. Fry, K. Slaw, N. F. Polizzi, Zero-shot design of drug-binding proteins via neural selection-expansion. bioRxiv, 2025.2004. 2022.649862 (2025).

40. S. A. Rettie et al., Cyclic peptide structure prediction and design using AlphaFold2. Nature Communications 16, 4730 (2025).

41. S. A. Rettie et al., Accurate de novo design of high-affinity protein-binding macrocycles using deep learning. Nature Chemical Biology, (2025).

42. X. S. Wang et al., A genetically encoded, phage-displayed cyclic-peptide library. Angewandte Chemie 131, 16051–16056 (2019).

43. H. C. Hayes, L. Y. P. Luk, Y. H. Tsai, Approaches for peptide and protein cyclisation. Org Biomol Chem 19, 3983–4001 (2021).

44. Q. Li, D. Daumiller, P. Bryant, RareFold: Structure prediction and design of proteins with noncanonical amino acids. bioRxiv, 2025.2005. 2019.654846 (2025).

45. Q. Li, E. N. Vlachos, P. Bryant, Design of linear and cyclic peptide binders from protein sequence information. Communications Chemistry 8, 211 (2025).

46. G. R. Lee et al., Small-molecule binding and sensing with a designed protein family. bioRxiv, 2023.2011. 2001.565201 (2023).

47. L. Pinheiro, C. Faustino, Therapeutic strategies targeting amyloid-β in Alzheimer’s disease. Current Alzheimer Research 16, 418–452 (2019).

48. S. Kumar, J. Walter, Phosphorylation of amyloid beta (Aβ) peptides–A trigger for formation of toxic aggregates in Alzheimer’s disease. Aging (Albany NY) 3, 803 (2011).

49. K. Wu et al., Design of intrinsically disordered region binding proteins. Science 389, eadr8063 (2025).

50. P. Li et al., FGFR2 promotes expression of PD-L1 in colorectal cancer via the JAK/STAT3 signaling pathway. The Journal of Immunology 202, 3065–3075 (2019).

51. S. Palakurthi et al., The combined effect of FGFR inhibition and PD-1 blockade promotes tumor-intrinsic induction of antitumor immunity. Cancer immunology research 7, 1457–1471 (2019).

52. X. Ye, R. A. Kumar, D. J. Patel, Molecular recognition in the bovine immunodeficiency virus Tat peptide-TAR RNA complex. Chemistry & Biology 2, 827–840 (1995).

53. A. Kubaney et al., RNA sequence design and protein–DNA specificity prediction with NA-MPNN. bioRxiv, 2025.2010.2003.679414 (2025).

54. L. Huang et al., A dual diffusion model enables 3D molecule generation and lead optimization based on target pockets. Nature Communications 15, 2657 (2024).

55. M. Li et al., Electron-density-informed effective and reliable de novo molecular design and optimization with ED2Mol. Nature Machine Intelligence 7, 1355–1368 (2025).

56. V. Le Guilloux, P. Schmidtke, P. Tuffery, Fpocket: an open source platform for ligand pocket detection. BMC Bioinformatics 10, 168 (2009).

57. M. van Kempen et al., Fast and accurate protein structure search with Foldseek. Nature Biotechnology 42, 243–246 (2024).

58. P. D. Adams et al., PHENIX: a comprehensive Python-based system for macromolecular structure solution. Acta Crystallogr D Biol Crystallogr 66, 213–221 (2010).

59. P. Emsley, K. Cowtan, Coot: model-building tools for molecular graphics. Acta Crystallogr D Biol Crystallogr 60, 2126–2132 (2004).

